# Subsequent Phases of Gene Expression Leading to Commitment and Differentiation in a Eukaryotic Cell

**DOI:** 10.64898/2026.11.24.768869

**Authors:** Wolfgang Marwan, Markus Haas, Monika Heiner

## Abstract

Cell differentiation requires the specific control of gene expression for each cell type. However, the mechanism by which robust developmental outcomes arise from inherently noisy single-cell dynamics remains unclear. To address this question, we performed a formal analysis of longitudinal time series during the commitment and differentiation of giant single cells. We tracked how the discrete expression states of individual genes changed relative to each other. Despite marked variability in timing and trajectories from cell to cell, we identified reproducible temporal motifs and quantified the strength of temporal correlations among early- and late-regulated genes. Notably, a core gene set exhibited the same relative temporal order in establishing stable or transiently stable expression states in most cells. This suggests that a well-defined gene expression program emerges amidst apparent stochasticity. These results support a model in which induction to differentiation proceeds via a cascade of partially concurrent, discrete genetic switching events that collectively produce gradual, continuous shifts in expression at the population level. This framework reconciles apparently stochastic single-cell behavior with reliable developmental outcomes and suggests general principles for the temporal organization of gene regulation during differentiation. Furthermore, it provides analytic tools that can be broadly applied to longitudinal single-cell datasets to design temporally precise interventions in development and disease.

## Introduction

The process by which eukaryotic cells differentiate into functionally specialized cell types involves the differential regulation of gene expression, which ultimately leads to the formation of stable, cell-type-specific gene expression patterns. This process involves intricate networks of biomolecules that interact in complex ways. The functional dynamics that determine cell fate remain elusive (Endres 2012; Rowland et al. 2012; Varusai et al. 2015; Vilar et al. 2003; Bornholdt and Kauffman 2019; Greulich et al. 2020; Huang 2011).

From a theoretical perspective, the Waddington landscape paradigm provides a foundation for formally analyzing these dynamics (Bornholdt and Kauffman 2019; Greulich et al. 2020; Huang et al. 2009; Marr et al. 2016). However, pseudo-time series, as are usually obtained from single-cell RNA sequencing (RNAseq), pose fundamental limits on the reconstruction of functional dynamics (Weinreb, 2018; Greulich, 2020). Here, we take an orthogonal approach, analyzing true longitudinal time series of differentiating giant *Physarum polycephalum* cells. This amoebozoan model eukaryote develops different cell types throughout its life cycle. This allows for the time-resolved analysis of fundamental phenomena in cell differentiation and developmental biology with defined phases of competence, induction, commitment, morphogenesis, meiosis, and game-togenesis (Sauer 1982; Sauer et al. 1969; Wick and Sauer 1982). Transcriptome analyses have shown that commitment and subsequent differentiation processes in *Physarum* giant cells are associated with extensive changes in gene expression patterns involving regulatory processes at transcriptional, translational, and post-translational levels (Barrantes et al. 2010; Barrantes et al. 2012; Glöckner and Marwan 2017; Martel et al. 1988; Putzer et al. 1984). Due to their natural mitotic and developmental synchrony, *Physarum* giant cells (also known as plasmodia) provide large amounts of homogeneous cell material that can be repeatedly sampled over time to create true longitudinal time series (Guttes and Guttes 1961, 1964; Hoffmann et al. 2012; Pretschner et al. 2021; Rätzel and Marwan 2015; Rusch et al. 1966; Sachsenmaier et al. 1972; Starostzik and Marwan 1995a; Walter et al. 2013).

Here, we analyze longitudinal time series of individual plasmodial *Physarum* cells measured after differentiation induction. Despite the significant variation in overall gene regulatory dynamics, we identified a number of genes, including transcription factors, that transitioned to a stable or metastable expression state in a consistent temporal pattern in nearly all of the examined cells. Our results are based on a new computational analysis method (Marwan et al. 2024), that can be applied to any longitudinal single-cell time series data. Major steps are described in the Results and Supplementary Methods and Results sections.

## Results

### Time Series Dataset

We used a gene expression dataset of competent giant cells (plasmodia of *Physarum polyce-phalum*), which were induced to sporulate by a short pulse of far-red light (Haas et al. 2026a, b; Pretschner et al. 2021). This dataset contains longitudinal time series of the expression levels of 112 genes in each cell with two samples taken from each cell at ten one-hour intervals following an inductive differentiation stimulus pulse (Fig. 1A; SI Figs. 1,2) as determined by quantitative RT-PCR (Marquardt et al. 2021). The homogeneity of the protoplasm volume, which mixes spontaneously through shuttle streaming and synchronizes naturally, was validated through biological replicates taken from each cell at each time point (Haas et al. 2026a; Pretschner et al. 2021).

**Figure 1.**
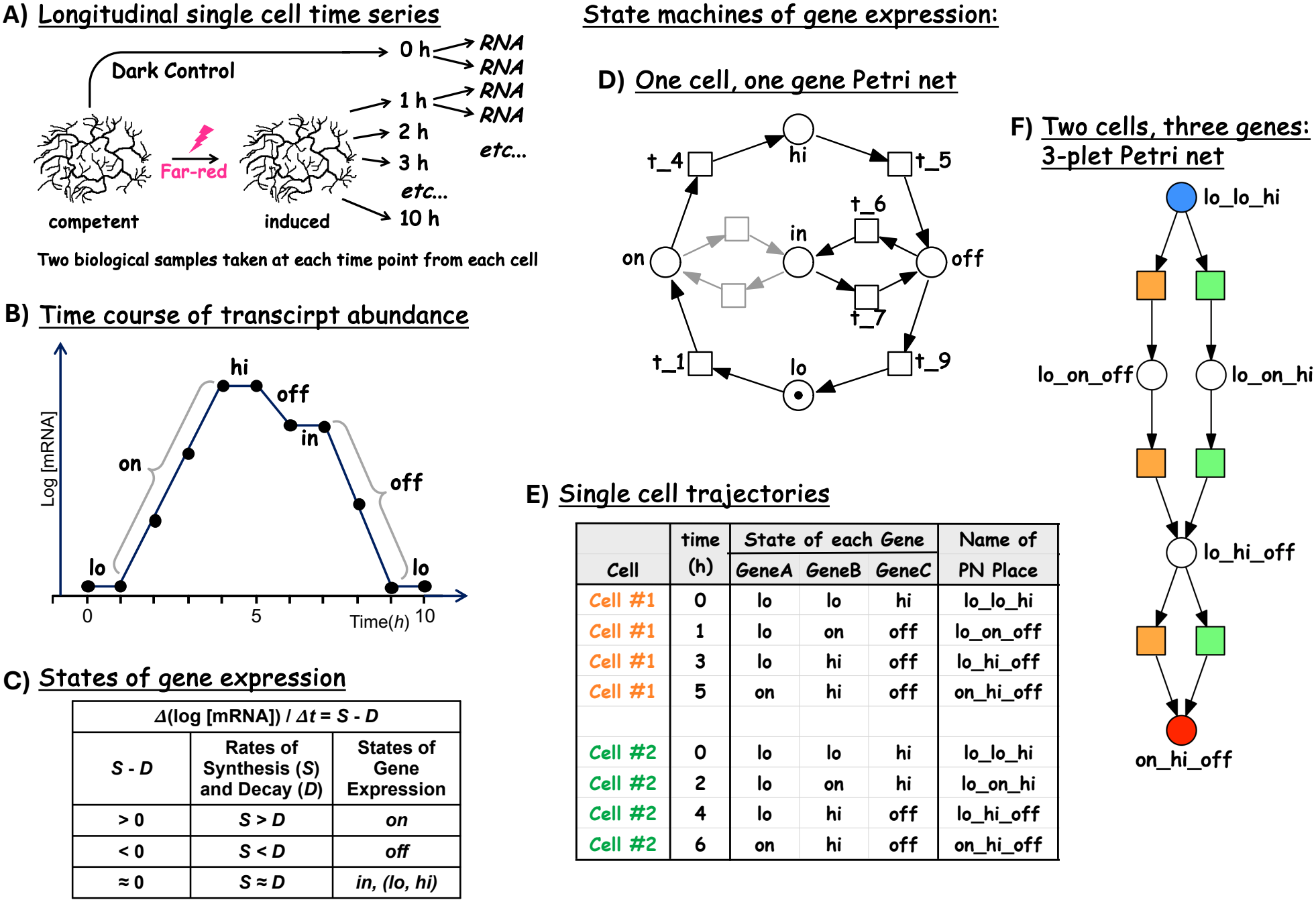
Longitudinal time series of gene expression in individual cells and analysis of changes in gene expression status. (A) Competent multinucleate giant cells of *P. polycephalum* are induced to sporulate by a far-red light pulse. Two samples of protoplasm are collected immediately before and at one-hour intervals after the stimulus to obtain a longitudinal time series from each cell with biological replicates from the same cell. Depending on whether the concentration of an mRNA increases (*on*), decreases (*off*) or remains at a low (*lo*), high (*hi*) or medium (*in*) level without changing significantly, a correspondingly named discrete state of gene expression is defined (B, C). The trajectory of a cell consists of a temporal sequence of these states. (D) A state machine can be constructed from the temporal sequence of expression states of a gene in a cell and represented in the form of a Petri net, in which the states are drawn as circles and the transitions between states as rectangles. In the time series shown in (B), no transits occurred between the states *on* and *in*, which is why the corresponding transitions, shown in grey, are missing from the state machine. (E) The expression states of multiple genes prevailing in a given cell at a given time can be combined into a global state. The trajectory of a cell consists of the temporal sequence of these aggregate states. (F) Following the principle shown in (D), a state machine can be reconstructed in the form of a Petri net from the trajectories of multiple cells. The net represents the behaviour of the cell population and the changes in the expression states of various genes relative to one another.

To compare the differential gene regulation of individual cells, we plotted the time course of all examined transcripts for each of the 16 light-induced cells and found considerable variability in kinetics from cell to cell (SI File 1). To provide an example of how much the time courses of different mRNAs vary relative to each other, we plotted the kinetics of transcripts from sets of randomly selected, differentially regulated genes in the same plot and found that the patterns varied between cells (SI Fig 3). Analysis of the time course of log[mRNA] values revealed sudden changes in the expression kinetics of many genes. Generally, the time until a next sudden change appeared to be linear or almost linear (SI File 1). Successive phases of increase or decrease were often interrupted by phases of variable length during which the concentration of a transcript did not change significantly.

### Discretization

We discretized the time course of mRNA concentrations into three states depending on whether the concentration increased (on), decreased (*off*), or remained constant (*in*, for “invariant”). Additionally, states in which the concentration of an mRNA did not change significantly were specified further if they stayed at a relatively low (*lo*) or high (*hi*) level (Fig. 1 B,C). States *lo* and *hi* are special *in* states that represent the extreme values of respective mRNA concentrations. This allowed us to determine whether a gene was up- or down-regulated overall over the course of the time series. Discretization was performed by sampling the curves’ time courses, whereby the algorithm detected and statistically evaluated sudden changes and assigned them to the aforementioned states (Fig. 1B; SI Fig, 4, see the Supplementary Methods for details). This resulted in gene expression trajectories as temporal sequences of changes in discrete expression states (*on, in, off*, etc.) for each gene and cell.

To compare the time course of expression of individual genes in different cells, we constructed state machines in the form of a Petri net. In these Petri nets, the state of gene expression in a cell is represented by a place, which is depicted as a circle. Transit between successive states is represented by a transition, which is depicted as a rectangle connected to a place by a directed arc (Fig. 1D). The current gene expression state of a cell is indicated by a token marking the corresponding place. This token moves through the network from place to place in the directions defined by the edges. Thus, the gene expression trajectory, in terms of the temporal sequence of expression states of one or more genes, can be represented in comparison to different cells (see Fig. 1 D-F). For a formal definition of these Petri nets and further details, see Supplementary Methods and Results (Appendix 1).

### The temporal dynamics of gene expression in different cells

To visualize the temporal dynamics of the expression states of a single gene in individual cells, Petri nets were generated and the transitions were color-coded according to the time at which the respective expression state changed (Fig. 2). While the expression states of many genes changed in the same temporal sequence in many cells, the timing of these changes varied from cell to cell. The varying complexity of the resulting single-gene Petri nets indicates correspondingly different patterns of regulation. Some genes changed state only once or twice after differentiation induction, resulting in a relatively simple Petri net structure (*morC*; Fig. 2C). Other genes underwent several state changes over time, often with bifurcations of main trajectories or recurring expression states (Fig. 2A). Other genes, such as the transcription factor *bzpJ* (Fig. 2B), had periods of the same state (e.g., *on*) separated by longer or shorter periods (*in* states) during which the concentration of mRNA did not change significantly. The complete set of single-gene Petri nets for all the differentially regulated genes analyzed is shown in SI File 2, corresponding to the plots of log mRNA concentration versus time in SI File 1. Overall, the relative frequency of switching events per time interval remained relatively uniform, decreasing by approximately 30% from the beginning to the end of the time series (Fig. 2).

**Figure 2.**
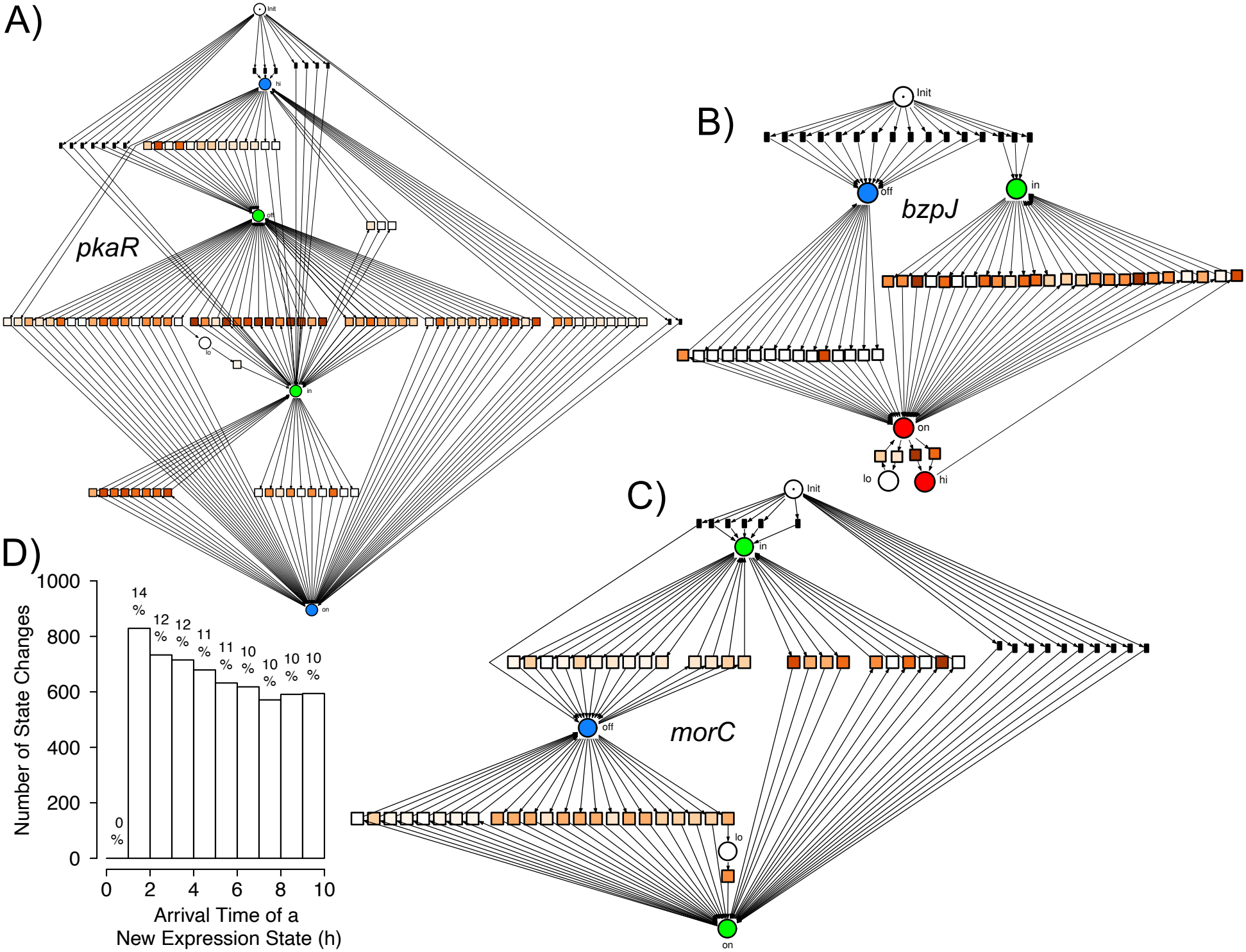
Time Course of Expression States for Individual Genes. Each Petri net illustrates a gene’s expression states over time in the analyzed cells. Transitions are color-coded according to when the corresponding transition occurred (early, light; late, dark). Places representing initial and/or final states are color-coded as well (blue for initial states, red for final states, and green for both). (D) Frequency of changes in the expression state of all genes over time.

### Correlated changes in gene expression states

Considering the varying complexity of regulatory patterns observed in single-gene Petri nets, we searched for genes that change state in a correlated manner, either simultaneously or in a specific temporal or partial sequence. Initially, we visually inspected a large number of Petri nets constructed for randomly selected combinations of three or four genes. We found Petri nets in which parallel transitions were arranged in a cascade-like manner. These parallel transitions defined main trajectories that groups of cells followed by undergoing the same temporal sequence of gene expression states (Fig. 3). We observed that the number of highly connected places and the number of parallel transitions between place pairs varied considerably depending on the gene combination under consideration (not shown). Highly connected places were not necessarily connected to each other by multiple parallel transitions but revealed two structural motifs that we refer to as hubs and diamonds (for a formal definition, see Appendix 1 in the Supplementary Methods and Results section).

**Figure 3.**
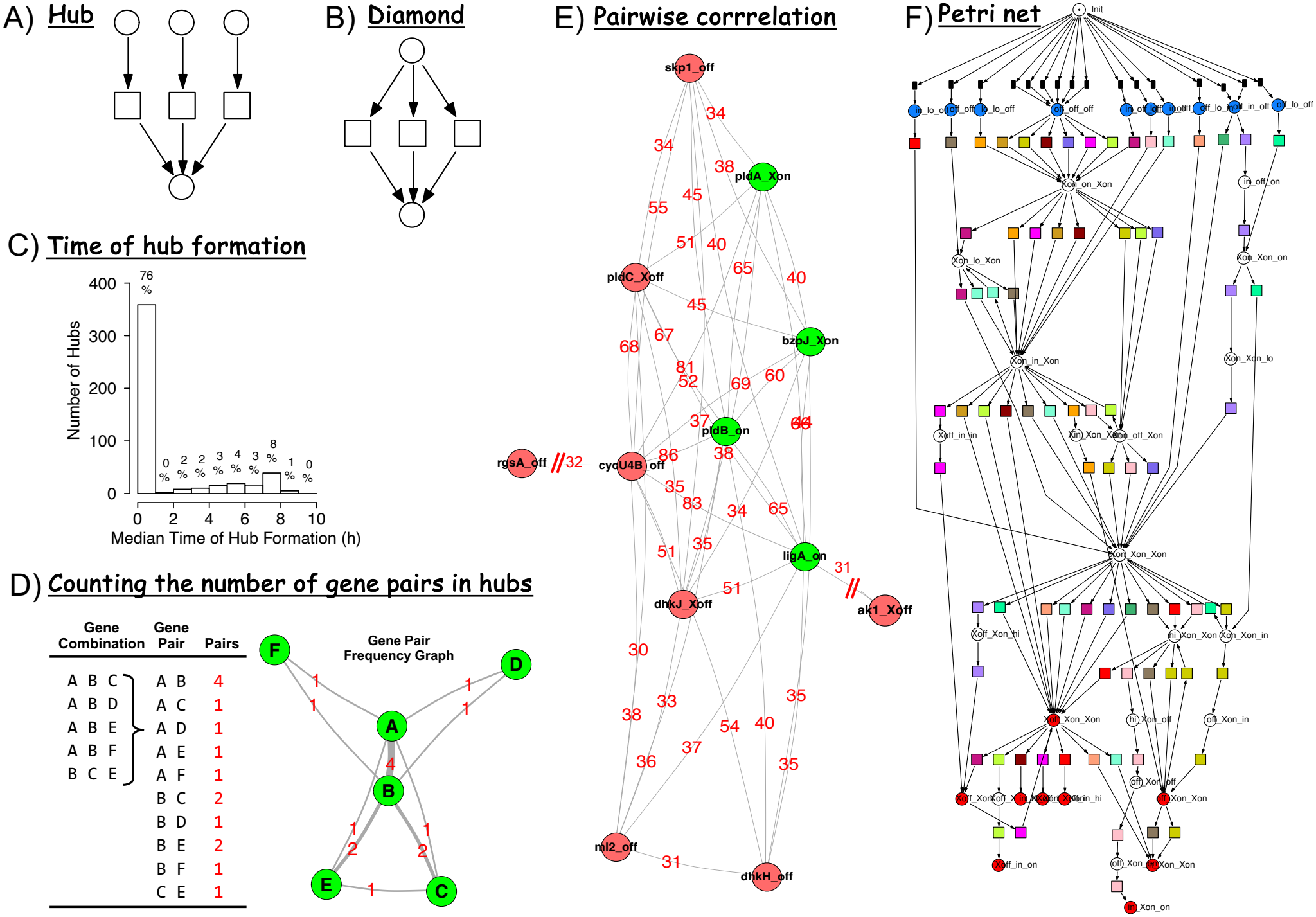
Analysis of Temporally Correlated Changes in Gene Expression Levels. Changes in gene expression states can give rise to hubs (A) and diamond-shaped (B) structural motifs in x-plet Petri nets. (C) Temporal distribution of hub formation over the time course of the time series. (D) The frequency with which individual genes contribute to hub formation is a measure of the strength of correlation in their differential regulation. (E) The strength of correlation among early differentially regulated genes in terms of gene pair frequency in hubs of 4-plet Petri nets is shown in a Fruchterman-Reingold plot. The stronger the correlation, the smaller the distance between two nodes. Red numbers indicate the number of hubs to which each gene pair contributed. Due to space limitations, the results obtained for extended states are shown. (E) Petri net constructed for the 3-plet of closely correlated genes *pldA, ligA*, and *bzpJ* indicating cascade-like regulation. The color of each transition is specific for the transits of an individual cell, highlighting single cell trajectories of gene expression.

A hub is a place with a defined minimum number of incoming edges originating from upstream transitions. Each edge is directly connected to a different pre-place (input place). A diamond, on the other hand, consists of a pair of places directly connected by a minimum number of parallel transitions in the same direction (Fig. 3 A,B). An important criterion for interpreting hubs and diamonds is the number of genes that change their expression state simultaneously when passing through the structural motif, as well as the number of cells involved in this change. For a given set of genes, hubs and diamonds reveal the previous states through which cells transitioned to reach a common state. Since hubs and diamonds occurred at different frequencies in Petri nets with randomly selected gene combinations, we used these structural motifs to algorithmically search for temporally correlated gene expression. To perform this analysis, we examined groups of four genes (4-plets) and created Petri nets for all possible combinations of the 72 most differentially regulated genes (SI File 1).

### Analysis of the Hubs

First, we examined how the number of hubs in these 4-plet Petri nets changed with the minimum number of incoming edges, as well as how many cells contributed to the formation of each hub. Presumably by chance, more than 80% of transitions involving four genes changing their expression status simultaneously emerged from the behavior of one individual cell. As expected, the number of transitions in which four genes simultaneously changed their expression status in a certain minimum number of cells decreased sharply with the specified number of cells. To avoid setting the minimum number of cells too strictly and to examine the variability of gene expression patterns from gene to gene and cell to cell, we searched for hubs in which the expression status of four genes changed simultaneously in at least six of the 16 analyzed cells.

Hubs occurred most frequently immediately after the light stimulus (Fig. 3C), suggesting, that these correlations emerge from a group of immediately responding genes. Due to the lack of strict coupling, these genes seem to be regulated in a concurrent manner. The scarcity of correlations of state changes of other genes during the subsequent period of the time series suggests a relatively high degree of concurrency in the regulation of light-induced gene expression and is possibly due to the relatively small number of genes considered in this study.

Structural analysis of the 4-plet Petri nets identified combinations of genes that generated a relatively large number of hubs. This phenomenon and its quantification are shown schematically in Fig. 3D using a 3-plet Petri net as an example. Two genes, A and B, are identified as changing their expression state simultaneously with particular frequency. If all genes forming hubs in multiple combinations were strictly correlated or co-regulated in expression, all transitions leading to corresponding hub places would have to be assigned to the same group of cells. However, the ID numbers of the cells indicate that the hubs, which emerged from different combinations of members of a subset of a gene set, were formed by disjunct subgroups of cells. Therefore, the state changes of the hub-forming genes were not strictly coupled. Certain gene pairs contributed much more frequently, i.e., with a greater probability, to the occurrence of a hub than others. This suggests that the response to the inductive far-red light pulse took the form of largely concurrent processes that were temporally correlated with a certain probability, but not strictly coupled to each other.

The strength of the temporal correlation with which genes changed their expression state was determined by the frequency with which each gene pair occurred in the hubs of the total number of 4-gene Petri nets considered. This frequency was represented using the Fruchterman-Rein-gold graph layout algorithm Fig. 3E). A Petri net constructed for a 3-plet of closely correlated genes *pldA, ligA*, and *bzpJ* indicates a partly correlated, cascade-like regulation (Fig. 3F).

### Analysis of Diamonds

We searched for diamonds, as defined above, in the entire set of 4-plet Petri nets constructed for all combinations of the 72 highly differentially regulated genes. Similar to the hubs, we characterized each diamond in terms of the number of contributing cells, the number of transitions involved, and the number of genes that changed their expression state. As with the hubs, the number of diamonds decreased with the increasing threshold set for the minimum number of transitions. In the vast majority of diamonds in 4-plet Petri nets with at least six parallel transitions, only one gene changed its expression state while the states of the other three genes remained unchanged. However, many of these genes had previously changed their expression state, revealing transiently constant expression states against which the respective change took place (not shown). This suggested a possible temporal sequence in the expression of certain genes.

### Temporal Sequence of Gene Expression

Given the high variability and concurrent nature of stimulus-induced gene regulation, the apparent scarcity of strict correlations, and motivated by the results of the Diamond analysis, we searched for genes that are regulated in a defined temporal order. Upon reviewing the plots that compare the time course of log mRNA concentration for each analyzed cell (see SI File 1), we noticed several genes whose log mRNA concentration increased or decreased by one to two orders of magnitude in a linear fashion. The exponential increase suggests positive feedback. Phases of increase or decrease began at different times after the inductive photostimulus and lasted for several hours. This suggests that a new expression state of the respective gene was established as a result of a switching event which remained relatively stable over time during the observation period, unlike the fluctuating expression states of other genes. To distinguish such states from merely transient changes, we introduced the concept of “extended states,” designated *Xon, Xoff*, and *Xin*, when a state persisted for a defined minimum time. For further analysis, we focused on the subset of genes that assumed a stable or metastable state (*Xon* or *Xoff*) in nearly all cells. Using a pairwise search, we identified genes that entered an extended state in the same temporal sequence in almost every cell, regardless of the chronological timing of the switching events relative to the stimulus pulse. We found that this regulation occurred from quite different preceding states.

The Dumbbell plot in Fig. 4 shows the timing of these genes entering an extended state for each of the light-stimulated cells in comparison. All of the listed genes entered an extended state in a statistically significant order (FDR ≤ 0.05) relative to at least one other gene in the group (for details regarding the computational methods, the statistical analyses, and for the results of additional controls, see Supplementary Methods and Results). Despite the high variability seen from cell to cell, the Dumbbell plot gives the impression of cascade-like gene regulation. Some genes were already in an extended state at the time of stimulation, though this state may have been unstable if the cell had not been stimulated. The temporal patterns of gene expression vary greatly from cell to cell and are clearly visible in a Dumbbell plot where the behavior of each cell is shown in a separate timeline (SI Fig. 8B,D; SI Fig. 13).

**Figure 4.**
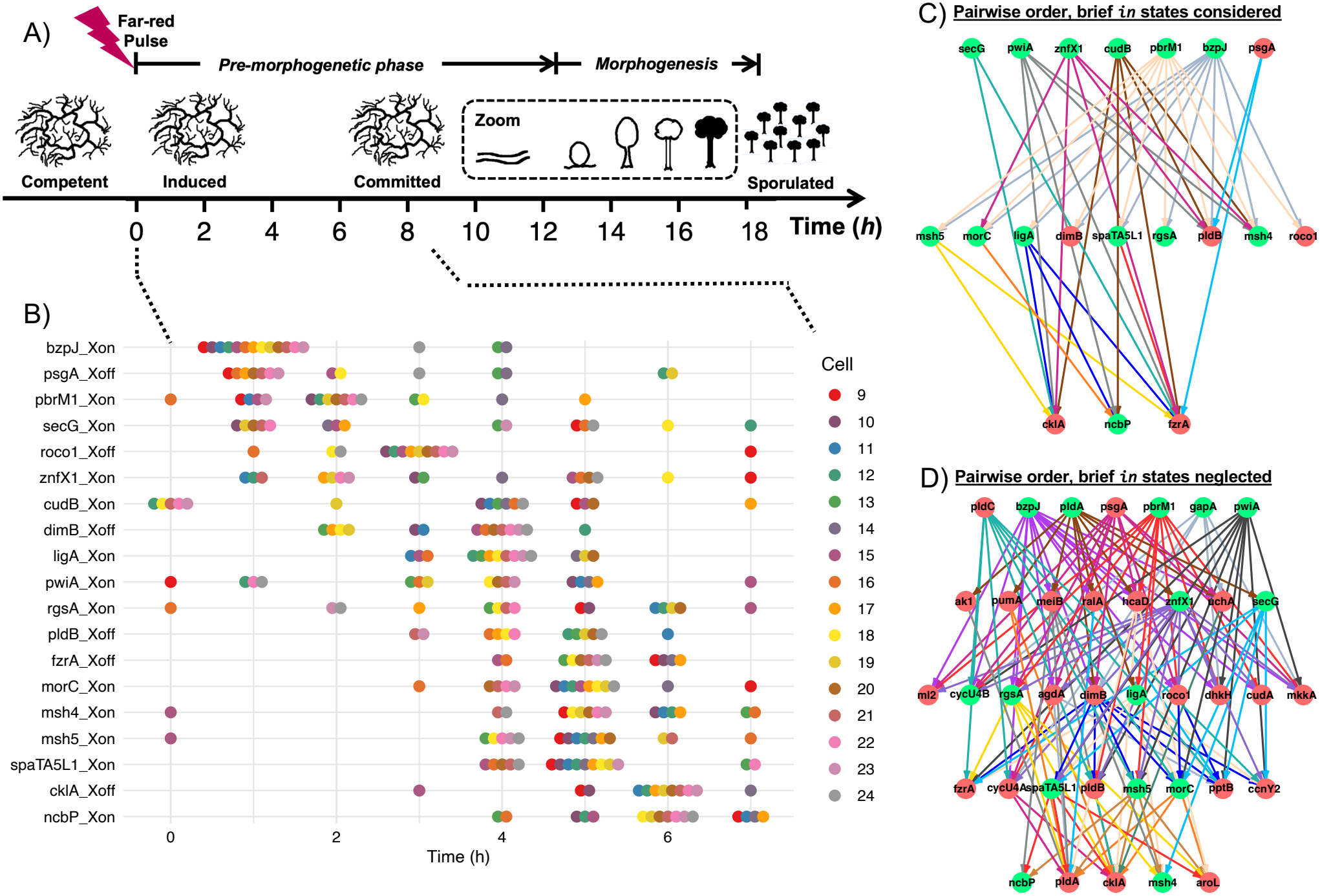
Time Course of Gene Expression Leading to Commitment and Differentiation. (A) The time course of induction, commitment, and differentiation during light-induced sporulation of *P. polycephalum*. The plasmodium is induced by a brief pulse of far-red light, followed by a pre-morphogenetic phase in which no visible change occurs. Commitment, or the irreversible determination toward differentiation, occurs approximately four to six hours after induction. Approximately 12 hours after the stimulus, the formation of fruiting bodies containing spores begins. The various stages of fruiting body formation are shown enlarged. (B) Temporal sequence of gene expression in the form of a Dumbbell plot. Each data point indicates the time at which a gene enters an extended state in a given cell. The data have a temporal resolution of one hour. Data points that correspond to the same time are plotted side by side. (C, D) The temporal order of gene expression is shown in the form of an iGraph plot using the Sugiyama layout. The arrows between nodes show the statistically significant temporal order (FDR ≤ 0.05) in which two genes enter an extended state. Panel C shows the plot taking brief interruptions in the rise or fall of transcript concentration into account. Panel D shows the plot with short interruptions in the rise or fall of transcript concentration disregarded. This results in many more genes exhibiting a statistically significant (FDR ≤ 0.05) temporal sequence in which they enter an extended state.

To quantitatively assess the existence of cascade-like regulation and check for possible concurrency, we grouped genes that were regulated in a statistically significant order to create a coherent graph (Fig. 4C). Each directed arc indicates a statistically significant (FDR ≤ 0.05) temporal sequence in which the two connected genes entered an extended state. Redundant arcs are not shown. Input nodes represent genes for which no significant temporal order could be determined relative to each other. The same applies to output nodes. A version of this figure in which the vertical position of the nodes corresponds to the average time at which the respective gene entered the extended state is shown in SI Fig. 14. Thus, despite the high variability in differential gene regulation, a group of genes exists that are regulated in the same temporal order in all or almost all cells. The most frequent gene ontology terms and superterms for these genes are related to signaling, response, differentiation, meiosis, and proliferation control (SI Fig. 16). Due to their relatively strict temporal control in relation to one another, these genes are ideal candidates for reference genes in further mechanistic analyses and network reconstruction.

As mentioned above, the temporal profile of transcript concentration for many genes is interrupted by *in* states, during which the mRNA concentration does not change significantly. The cause of these *in* states is unclear. However, we do not assume that this phenomenon is due to measurement artifacts since the curve segments before and after an *in* state often appear shifted parallel along the time axis (SI File 1). Further and more precise measurements are required to definitively clarify whether *in* states are a physiological phenomenon. In any case, the effect of *in* states on the emergence of extended states is interesting, especially if the *in* states are physiologically relevant and mechanistically grounded. Therefore, we removed brief *in* states lasting no more than one hour from the existing dataset if they occurred between two *on* or two *off* states. This converted sequences *on* -> *in* -> *on* to *on* and *off* - > *in* -> *off* to *off*. In the edited dataset, we found approximately twice as many genes entering into an extended state as in the original dataset (36/19 = 1.89) for a time window of *Δt*_*w*_=0h (see Supplementary Methods and Results and SI Fig. 9). Consequently, the number of gene pairs entering an extended state in a significant temporal order (FDR ≤ 0.05) increased from 41 to 205. The largest number of genes that transitioned to an extended state in direct temporal order increased from three to five when the brief *in* states were removed (Fig. 4D; SI Fig. 14; SI Table 2). Further structural analyses using Petri nets suggest that the appearance of extended states reflects essential processes in the orchestration of cell differentiation (SI Fig. 15). These analyses also suggest that removing brief *in* states, regardless of their physiological relevance, provides a more comprehensive view of the actual events occurring within the cell (for details see Supplementary Methods and Results).

## Discussion

We analyzed true longitudinal time series of gene expression using a newly developed computational approach. These time series were recorded from individual multinucleate giant cells of *Physarum polycephalum* that were differentiating in response to phytochrome photoactivation (Lamparter and Marwan 2001; Starostzik and Marwan 1995b).

We discretized the time course of the x-fold change in mRNA concentration of 72 strongly light-regulated genes and determined discrete states of gene expression indicating whether the mRNA concentration increased, remained constant, or decreased. Building on an earlier approach (Rätzel et al. 2020; Werthmann and Marwan 2017), we constructed state machines from the temporal sequences of these states (single-cell trajectories) and represented them as Petri net graphs. These graphs comparatively depict the behavior of the analyzed cells for a small set of x genes, called an “x-plet”. For the genes under consideration, the paths that a token takes through a Petri net can be understood as a discrete sequence of gene expression patterns corresponding to paths through the Waddington landscape (Pretschner et al. 2021; Rätzel et al. 2020; Werthmann and Marwan 2017). The structures of the Petri nets exhibited significant variability in the temporal patterns of light-induced gene expression from cell to cell and from gene to gene. We found that certain gene combinations generate Petri nets with a cascade-like appearance in Sugiyama layout (Sugiyama et al. 1981). These nets have places representing successive temporal states that are connected by subsequent sets of parallel transitions. This is due to a certain number of cells changing the expression state of the considered genes in the same temporal order. A systematic search for two structural motifs — hubs and diamonds — revealed genes that switched on or off in a correlated manner with a relatively high probability, despite extensive concurrency in gene regulation. However, this correlation was not caused by the differential regulation being strictly coupled within individual cells because the strong temporal correlations were generated by non-disjunct subgroups of cells. We identified a group of genes whose expression state changed just one hour after the inductive photostimulus. This occurred well before the cells committed to differentiation by crossing the point of no return (Hoffmann et al. 2012; Starostzik and Marwan 1995a). Despite the pronounced variability and concurrency in changing expression states, and even though the observed correlations did not result from a strict coupling of differential gene regulation within a single cell, the results suggest that the response is structured to at least some extent.

Due to the relatively narrow and reproducible time window extending from light induction to the onset of nodulation and subsequent fruiting body formation (Fig. 4a; (Hoffmann et al. 2012)), we searched for a “fixed program” of differential gene regulation. We found a group of genes that had entered an extended state, which remained stable for several hours in all or almost all of the analyzed cells, depending of the gene considered. The sustained increase or decrease in mRNA concentration is presumably the result of a genetic switch. Despite the large variability in the timing of gene expression from cell to cell, some genes were regulated in a strict temporal sequence in almost all cells. These genes could be arranged in a corresponding cascade of gene expression (Fig. 4), indicating a structured response to the photostimulus.

This structured response was notably only seen in the longitudinal single cell true time series and disappeared when cell IDs were randomly shuffled to create pseudo-cell time series (see Supplementary Methods and Results for details). Thus, the gradual and continuous change in gene expression that occurs during differentiation, from induction through commitment and up to the onset of morphogenesis, is the result of a series of switch-like changes in gene expression states. These switching events occur concurrently and in a well-defined partial order for a set of genes in almost all of the examined cells. Switching to an extended *on* state (*Xon*) results in a sustained, often exponential, increase, which suggests positive feedback. Conversely, switching to an extended *off* state (*Xoff*) manifests as an exponential decrease in mRNA concentration. The well-defined temporal order of controlling a specific set of genes can be queried for mechanistic analyses and provides a scaffold for network reconstruction.

The edges of the graph in Fig. 4C,D represent statistically significant temporal sequences of regulation, rather than causal regulatory influences. Without additional information, causal regulatory interactions cannot be inferred from temporal correlations. Changes in the time course of gene expression patterns in differentiation mutants (Rätzel et al. 2013) however can help reconstruct regulatory interactions. In addition, analyzing mutants is helpful and necessary because it is unlikely that all regulators involved in controlling cell differentiation can be identified through differential gene expression alone.

*P. polycephalum* is an amoebozoan that shares a last common ancestor with opisthokonts. While this precludes direct transfer of the results to medically relevant mammalian cells, it allows for evolutionary considerations that may be helpful in identifying universally valid, evolutionarily conserved regulatory mechanisms (Bussey et al. 2017; Sebe-Pedros et al. 2025). A key question emerges when comparing gene regulation in *Physarum* with that of other organisms, including mammalian cells: To what extent has the multinucleate organization of the plasmodium, over the course of evolution, given rise to genes or gene regulation that differ from the typical gene regulation found in all other eukaryotes? The genome and transcriptome of *Physarum* reveal a standard set of eukaryotic genes and expression patterns (Barrantes et al. 2010; Glöckner and Marwan 2017; Schaap et al. 2016). However, are the mechanisms by which gene expression is regulated in a multinucleate cell fundamentally different from those in a mononucleate cell? The life cycle of *P. polycephalum* consists of several mononucleate cell types, such as spores, amoebae, flagellates, and zygotes, as well as the multinucleate plasmodium. Thus, the genome can give rise to both mononucleate and multinucleate cells. Conversely, multinucleate cell types (syncytia and coenocytia) are found in plants, fungi, animals, and various human tissues. Some of these cell types can differentiate back into mononucleate cells. These findings suggest that the ability of an organism to form mononucleate and multinucleate cells, as well as the ability to switch between these cell forms, is deeply rooted in the evolution of eukaryotes. Thus, *P. polycephalum* represents a prototypical eukaryote in this respect.

A fundamental difference compared to most mammalian cells is related to the dynamics of gene expression and other biophysical aspects. As would be expected for mammalian cells due to substantial heterogeneity and variability in gene expression (Junker and van Oudenaarden 2014), we observed significant variations in the gene expression trajectories of individual *Physarum* cells. However, the gene expression dynamics of a multinucleate giant cell may differ considerably from those of a mammalian cell with a single nucleus. Most mammalian cell types have a small cytoplasmic volume and a single nucleus. During the G0 phase many genes are present in just two copies in a relatively small cytoplasmic volume which promotes stochastic fluctuations in gene expression. In the multinucleate *Physarum* plasmodium, the continuous mixing of the large cytoplasmic volume with the cell nuclei suspended therein leads to a natural synchronization of the cell nuclei (Guttes and Guttes 1961, 1964), and gene expression (Walter et al. 2013), as the suspended nuclei are exposed to a uniform concentration of cytoplasmic factors prevailing in the cell. Natural cytoplasmic mixing presumably buffers and averages out stochasticity in gene expression and transcriptional bursts, which occur in single-nucleated mammalian cells with typically small volumes. Dynamic regulatory phenomena based on nonlinear mechanisms may be more easily identified in *Physarum* because they are not obscured by stochastic fluctuations when analyzing macroscopic protoplasm samples. This could facilitate identifying causal dependencies within the regulatory network and help with network reconstruction.

## Acknowledgement

We thank Dr. George Assaf for implementing several options for a script-based batch processing of Petri net files.

## Availability of Material

All data, software, and supplementary information is available at https://doi.org/10.5281/zenodo.23954304.

## Suppementary Information

### Computational Methods

#### Time Series of Differential Gene Expression

The time series used in this study are derived from two published datasets (Haas et al. 2026a,b; Pretschner et al. 2021), which are described briefly below. The dataset consists of longitudinal time series from sixteen light-stimulated cells and eight unstimulated cells that served as dark controls. These controls demonstrate the effect of irradiation on gene expression. Stimulation was performed using a 15-minute, saturating far-red light pulse. All exposed cells sporulated in response to the stimulus, while none of the unexposed cells in the dark control sporulated.

Two protoplasm samples were taken from the stimulated cells at each time point: one sample was taken before irradiation (at 0 hours), and the other samples were taken at one-hour intervals after irradiation. This process was repeated to obtain a longitudinal time series of each single plasmodial cell ranging from 0 to 10 hours. Sampling was performed at different, relatively distant sites of the plasmodium to check for homogeneity of gene expression at each time point. Sampling from the unexposed cells was performed in the same manner.

Total RNA was isolated from each sample, and transcript concentrations of 112 genes were determined using independently performed GeXP RT-PCR assays (Marquardt et al. 2021; Haas et al. 2026a,b; Pretschner et al. 2021). Two technical replicates were obtained from each of the two biological replicates, corresponding to four values for each mRNA analyzed at each time point and cell. All samples were analyzed using three GeXP RT-PCR primer sets to ensure comparable relative transcript levels and temporal changes of the 112 genes in all samples (Haas et al. 2026; Pretschner et al. 2021). The mean values of the four samples at one time point and the standard errors of the means (SEM) of the irradiated cells were used as the basis for this study. The mean values were normalized to make the genes under consideration directly comparable in terms of x-fold changes in their transcript concentrations. To normalize the data, the values of a given gene’s transcripts across the entire dataset were divided by the geometric mean of all values for that gene. Mathematically, the logarithms of the individual values were averaged, and the resulting mean value from all the data for a given gene was subtracted from the logarithms of that gene’s individual values.

The resulting dataset is displayed in SI Figures 1,2. SI Fig. 1 shows the expression patterns of the eight dark-control cells compared to the sixteen light-stimulated cells. Each section of the heatmap represents the time series and hence the temporal expression pattern of a single cell. The left part of the figure enlarges sections of the time series from a non-stimulated, non-sporulating cell and a light-stimulated, sporulating cell, demonstrating the effect of light stimulation. SI Fig. 2 shows the entire dataset from SI Fig. 1 clustered using the hclust algorithm (Murtagh 1985) to highlight differentially regulated genes in the form of corresponding clusters. This representation solely serves to visualize the data. The functional annotation of the analyzed genes is provided as SI File 3.

#### Discretization of the time series

The four concentration values obtained for each time point and analyzed mRNA were averaged, log-transformed, and plotted against time to obtain the time courses of mRNA concentrations for the genes of interest in the form of kinetic curves for each cell, as described above. SI Fig. 3 shows a comparison of the expression kinetics of three genes from 16 cells, providing an arbitrary qualitative example of varying behavior from cell to cell. The kinetic curves of all gene transcripts can be found in SI File 1. We scanned the longitudinal time series of individual cells for potential changes in gene expression status using a moving double arrow that detects sudden changes in the time course of log[mRNA] concentration. This algorithm was originally developed to detect reversal points in the swimming direction of *Halobacterium* cells in response to a light stimulus (Marwan and Oesterhelt 1990). The double arrow scans the time course of the mRNA concentration point by point, evaluating three consecutive time points in each step. One arrowhead points to the preceding time point, and the other points to the time point immediately following the current one (SI Fig. 4). Abrupt changes (discontinuities) in the kinetic curve result in a small angle *β* formed by the two arms of the double arrow.

**SI Figure 1.**
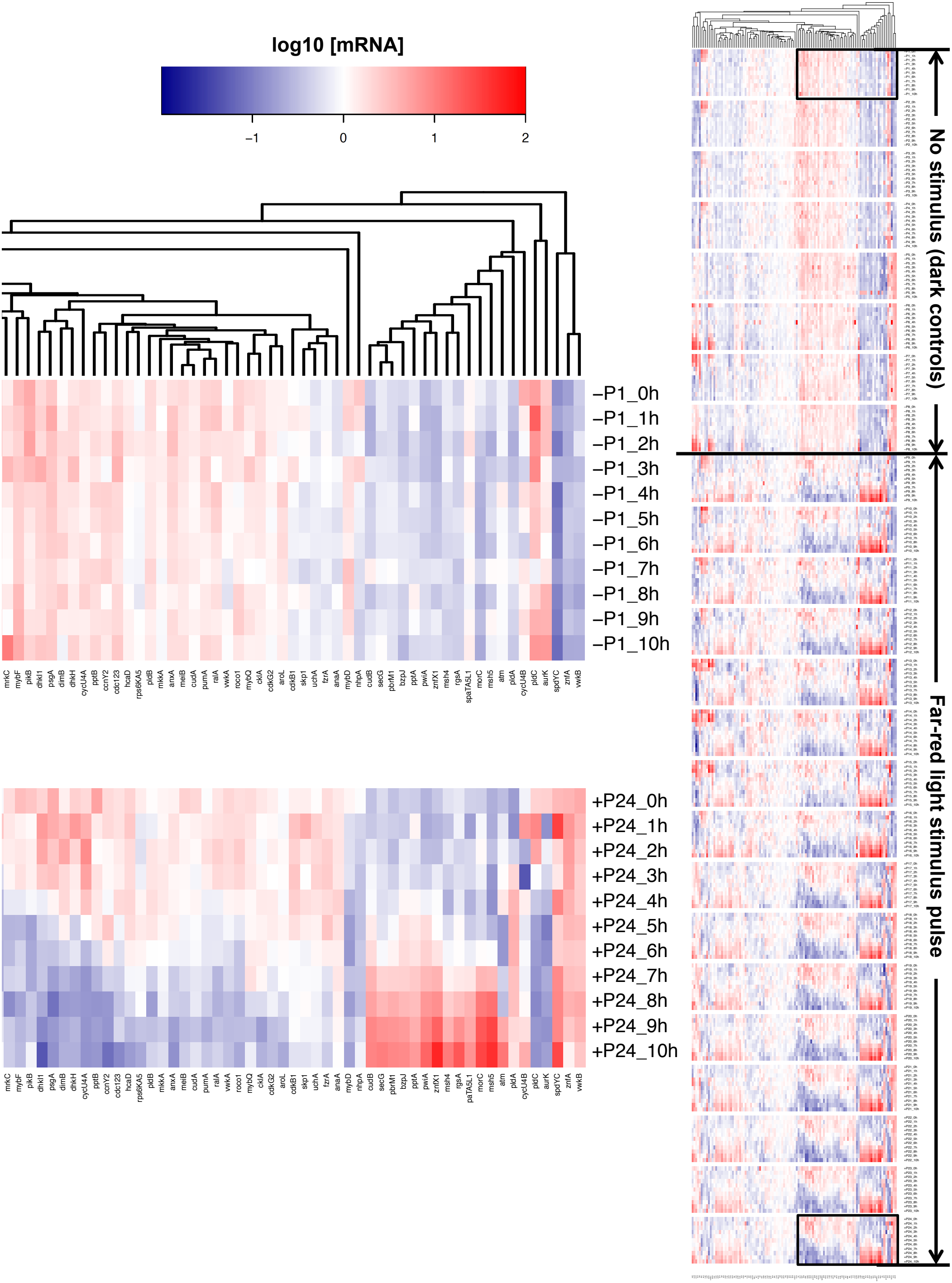
Longitudinal time series of stimulus-induced gene regulation in individual giant cells (plasmodia) of *Physarum polycephalum*. The heatmap on the right shows how the cells responded to a 15-minute far-red stimulus pulse (cells P9–P24) compared to the unstimulated dark control (cells P1–P8). All of the stimulated cells sporulated (+) in response to the stimulus, whereas all unstimulated cells did not (-). Each section of the heatmap displays the time course of log_10_ [mRNA] values of the analyzed genes in a single cell ranging from 0 to 10 hours. The 0 h value for the stimulated cells was recorded immediately before the stimulus pulse was applied. The left side of the figure shows enlarged datasets of an unstimulated cell (top) and a cell stimulated with a far-red light pulse (bottom). The data set shown was taken from (Haas et al. 2026a,b; Pretschner et al. 2021).

The algorithm identifies a break point as a potential change in gene expression status when angle *β* falls below a heuristically set threshold. Thus, the time course of each mRNA (log[mRNA] vs. time) was decomposed into a sequence of curve segments connected by break points. The data points of each curve segment, including the flanking break points, are fitted to a straight line via linear regression. The slope of the regression line corresponds to the log-fold change in mRNA concentration per unit of time, which is provided as an intermediate result (see SI Fig. 5).

A change in gene expression status following a break point is detected when the error bars (standard error of the mean [SEM] values) of the current break point do not overlap with those of the subsequent break point (see SI Fig. 4,5). Although the more stringent t-test criterion for testing the significance of the difference in mRNA concentration between two consecutive break points was implemented as an option, it was not applied since it often fails to detect obvious, continuous changes in gene expression status over a longer period of time.

Phases in which the mRNA concentration does not change significantly are classified as *in*. In Five-State Mode, an *in* state is classified as *hi* if the mRNA concentration of the first data point of the corresponding curve segment is at least 90% of the curve’s total concentration range and as *lo* if the mRNA concentration is no more than 10% of the total concentration range.

**SI Figure 2.**
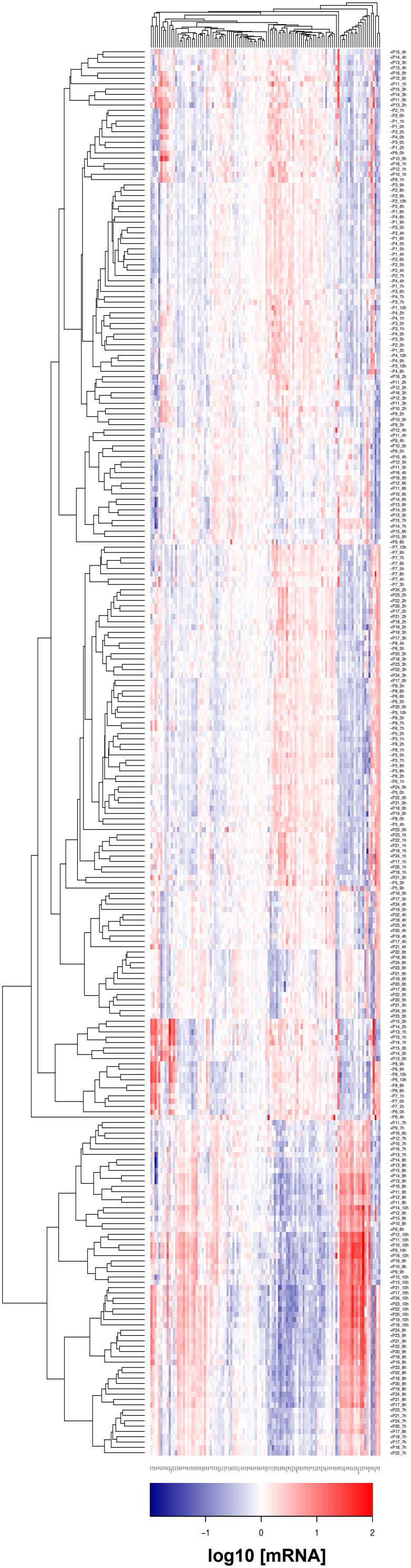
Time series data as in SI Figure 1, clusterd using the hclust algorithm.

**SI Figure 3.**
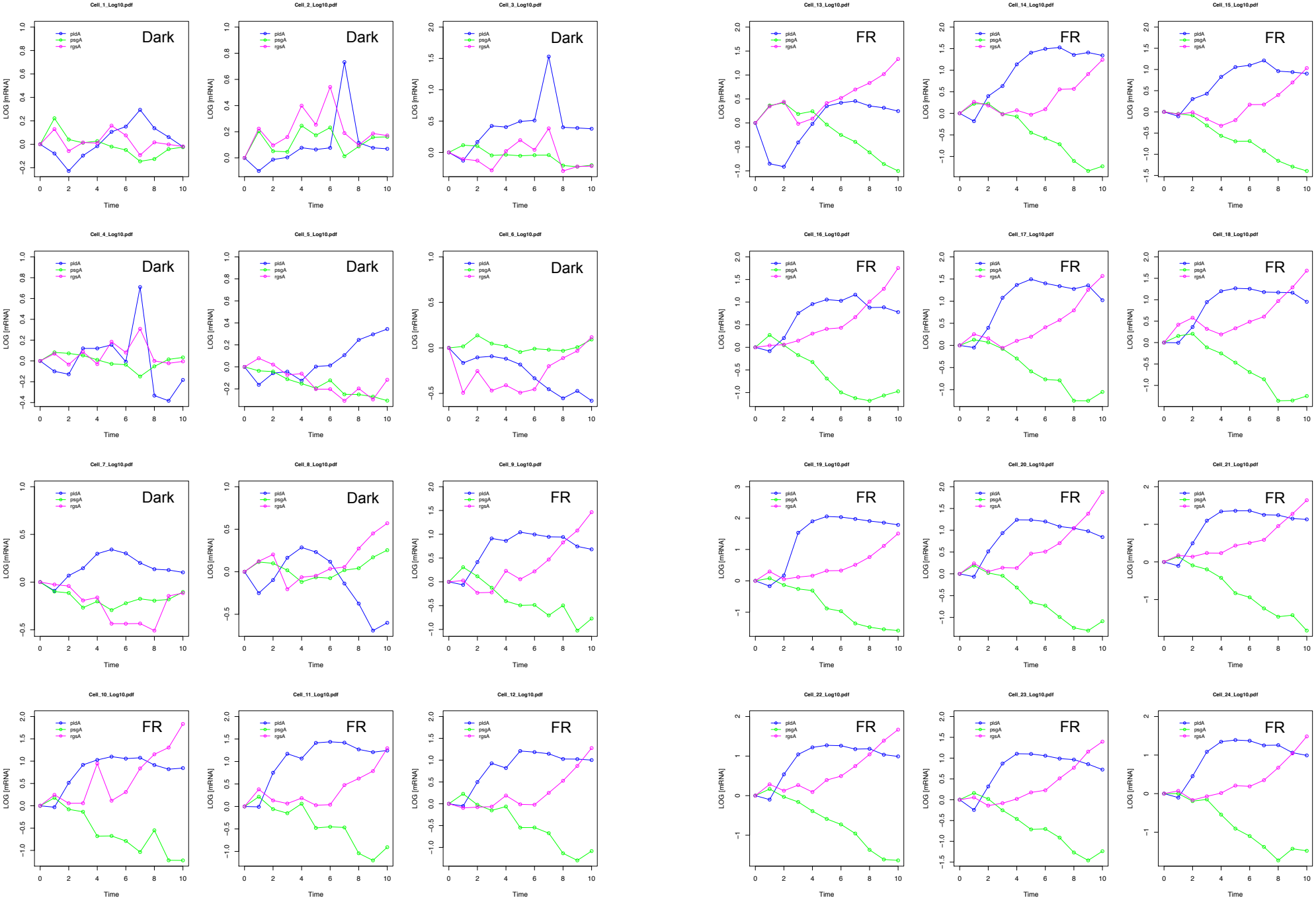
Gene expression kinetics of three selected genes (*pldA, psgA, rgsA*) in individual plasmodial cells. Each panel displays the expression kinetics of a different cell. Dark, dark control; FR, far-red stimulated.

The Check Backwards option retrospectively verifies whether consecutive phases of the same state (e.g., *on* or *off*) between potential break points are separated by an *in* state, during which the mRNA concentration does not change significantly. Without this check, the corresponding switching events (*on* -> *in* -> *on* or *off* ->*in* -> *off*) would go undetected in rare cases.

In the Check Backwards query, the system first checks if the error bars of the current break point and the potential reversal point overlap with the error bars of the preceding data point in the time series. If so, it checks if the error bars of the preceding data point overlap with the error bars of the preceding break point. If so, the state between the two points is set to *in*, as there has been no significant change in the expression state yet. Otherwise, the state of the preceding reversal point is retained, and a constant state (*in*) is inserted.

The intermediate results of the discretization calculation steps are documented in a series of tables and exported as a text file for each kinetic curve (see SI Fig. 5). To evaluate the functionality of the algorithm, the temporal concentration profile (log[mRNA] versus time) of each transcript can be plotted for each cell separately, and the cos# value, as well as the gene expression state determined by the algorithm, can be displayed next to the respective data point (see SI Fig. 4). Additionally, error bars (standard error of the mean) can be plotted to evaluate the significance of potential changes in expression status indicated by the double arrow. The result of the t-test evaluating the significance of concentration changes between two consecutive break points can also be displayed in the plot, if desired.

Visual inspection of the discretization results for hundreds of curves showed that the algorithm detects changes in gene expression status with high accuracy. Further improvement of the algorithm or the parameter settings did not appear necessary or useful for analyzing the existing dataset.

The final result of the discretization process is a so-called Node Table (SI Table 1A), which lists the successive expression states of each analyzed gene at every time point for each cell. This Node Table is the basis for constructing the Petri nets. Depending on whether the 3-state or 5-state model is used, the Node Table can be obtained for three (*in, off, on*) or five (*hi, in, lo, off, on*) states.

**SI Figure 4.**
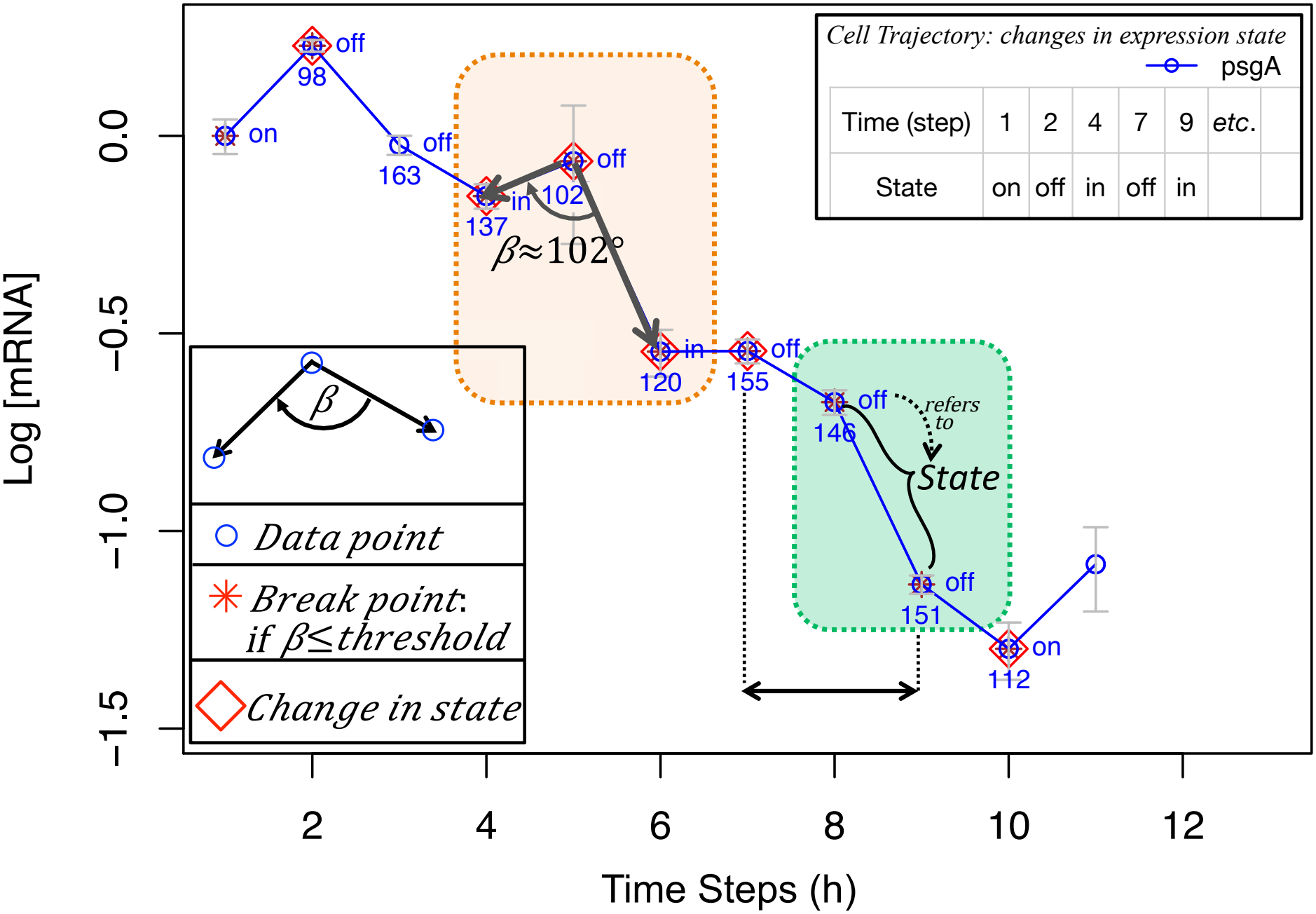
Scanning the time course of mRNA concentration to determine discrete states of gene expression using the double-arrow algorithm. The kinetics of each mRNA are scanned from one time point to the next using a double arrow. One tip of the double arrow points to the time point immediately preceding the current one, and the other tip points to the time point immediately following it. The two arms of the double arrow originate from the current time point. If the angle *β* formed by the two arrows falls below a threshold value (orange box), a break point of the curve—and thus a potential change in gene expression state—is detected, as described in the Computational Methods section. Break points detected by the double arrow are marked with a star. Those break points identified by the algorithm as changes in gene expression status based on additional criteria are marked with a rhombus (see the legend for SI Fig. 5). The angle *β* formed by both arrows is indicated below the respective data point. The gene expression state (*on, in, off*, etc.) determined by the algorithm and displayed to the right of each data point refers to the curve segment until the subsequent data point (green box). By incrementally scanning the kinetics, a single-cell trajectory is generated for the gene under consideration, in the form of a sequence of states and the corresponding time points at which the subsequent state begins. This is illustrated in the inset at the top right. The time course of each mRNA concentration in the dataset and the resulting state assignment can be output in the shown format.

#### Extended State Node Table

States that lasted for at least three hours were classified as extended states and stored in an Extended State Node Table. This table was derived by analyzing a previously generated Node Table. Before determining the extended states, short *in* states that occurred between two identical states (*on* -> *in* -> *on* or *off* -> *in* -> *off*) can optionally be removed from the original Node Table if they did not last longer than a user-defined maximum duration (in this study, one hour) (SI Figs. 6,7). In test mode, the program outputs a log of the computational steps and intermediate results for each gene and cell. This allows verification of the correct operation of the algorithm in detecting extended states.

**SI Figure 5.**
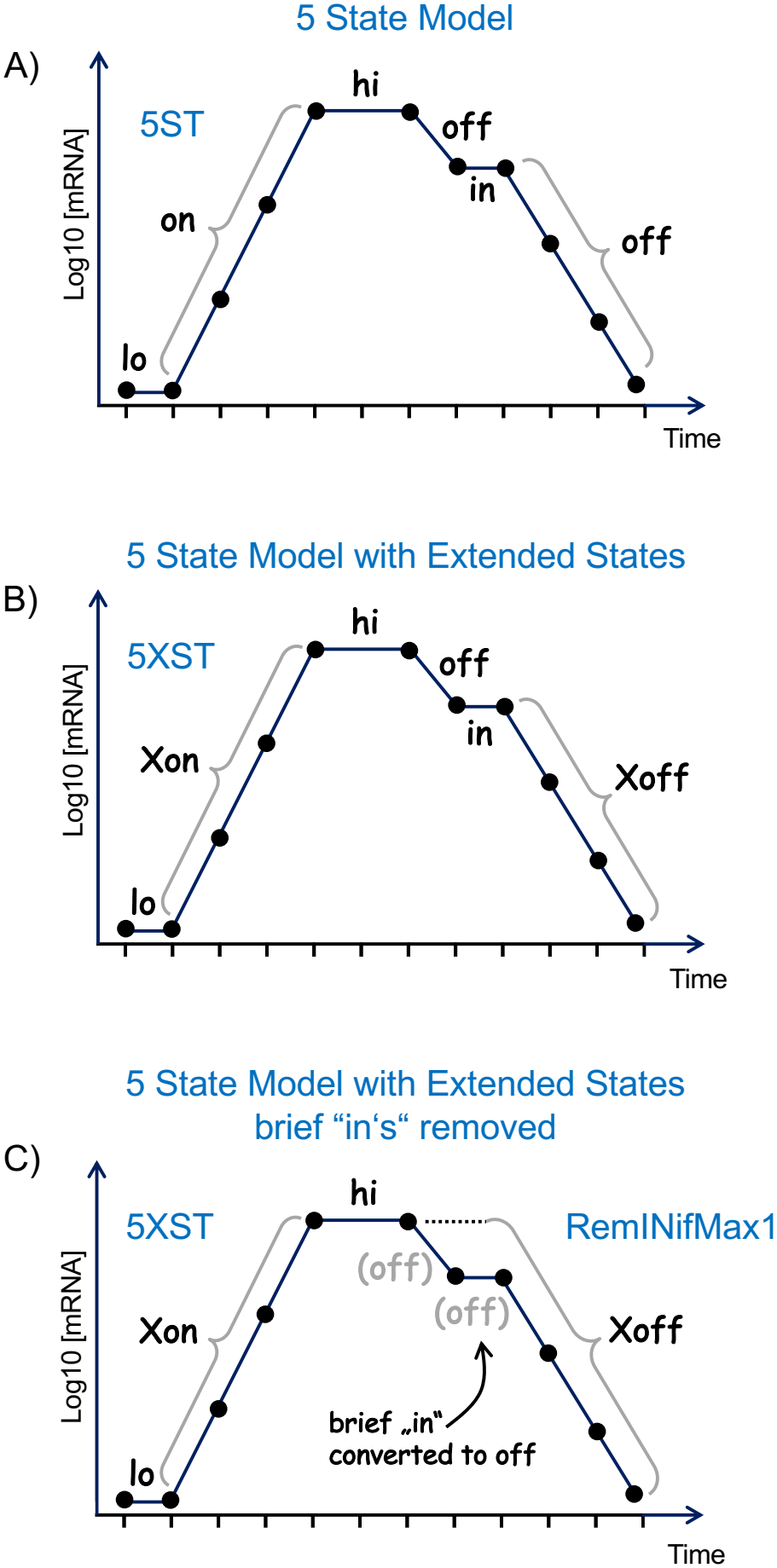
Intermediate results of the computational steps, as output during the determination of expression states using the double-arrow algorithm. The figure refers to the kinetics of two genes, *anxA* and *psgA*, from cell P9. Here, we focus on the intermediate results for *psgA*, which are highlighted in color. Using the double-arrow algorithm, break points in the time course of the logarithm of the mRNA concentration are determined, as shown in SI Fig. 4. The kinetics are thereby divided into a sequence of quasi-linear curve segments ((A); List of Curves). Each curve segment (segment 1 through 8) in the example shown in panel A and is displayed in a different color. Linear regressions are performed on the data points of each curve segment. The slope of each curve segment, as obtained by linear regression, corresponds to the x-fold change in relative mRNA concentration. B) The List of Changes lists the consecutive curve segments defined by break points, along with their start times and the parameter values obtained by linear regression as taken from (A). In the Changes with Raw States table (C), discrete expression states are assigned. To do so, the system checks if the error bars of the curve points in adjacent segments overlap. If they do not overlap (*ErrBarOverlap* yields FALSE), an expression state is assigned based on the slope of the curve segment (*on* for a positive slope, *off* for a negative slope). If the error bars overlap, the expression state associated with this curve segment is set to *const* regardless of the calculated slope, because the mRNA concentration has not changed significantly compared to the preceding data point. The *pvalues* column lists the results of the Student’s t-test. The values listed in the *x_fold_Ch_Thr* column for a user-defined concentration-dependent x-fold change threshold were evaluated as a possible criterion for a state change in an early phase of this study but finally not considered in the analyses. In the Changes with States table (D), the *on* and *off* states from Table (C) are carried over unchanged. However, the *const* states are specified depending on whether the 3-state or 5-state model is used. For the three-state model, all *const* states are set to *in*. For the 5-state model, *const* states where the mRNA concentration is ≤10% or ≥90% of the concentration range over which the entire curve extends are set to *lo* or *hi*, respectively; the remaining *const* states are set to *in*. These computational steps result in a trajectory (see the inset in SI Fig. 4) which consists of the successive expression states of a particular gene in a particular cell, assigned to the time points at which each state occurred.

**SI Figure 6.**
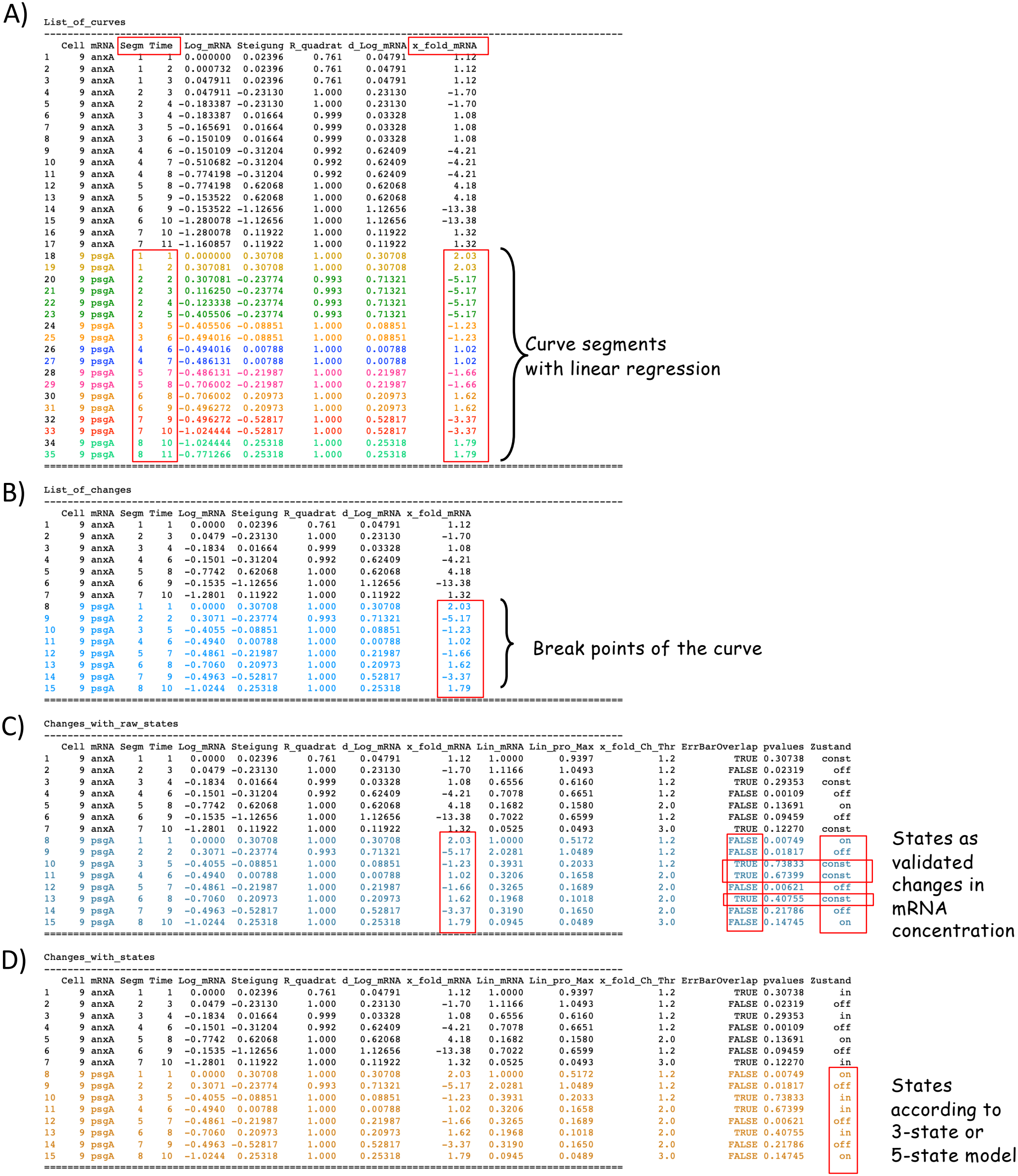
State models of gene expression. (A) Sudden changes in the time course of the logarithmic abundance of a gene-specific mRNA are analyzed using the double-arrow algorithm to determine a sequence of discrete gene expression states, as described in SI Figures 4 and 5. Depending on whether the mRNA concentration increases (*on*), decreases (*off*), or remains at a low (*lo*), high (*hi*), or medium (*in*) level, a corresponding state of the five-state model is defined. The trajectory of a cell consists of the temporal sequence of states or state changes for each analyzed gene. (B) The Extended State Model defines an Extended State when a state remains unchanged for an extended period, e.g., over three hours, as shown in the example. This allows one to determine whether and at what point a gene enters a stable or meta-stable expression state. (C) To examine the effects of brief interruptions of otherwise continuous states on the results obtained, short-lived *in* states located between two identical states can be disregarded. Sequences of *on* -> *in* -> *on* or *off* -> *in* -> *off* then turn into *on* or *off*, respectively, or, if long enough, into *Xon* or *Xoff* in the Extended State Model. As expected, removing short *in* states lasting no more than one hour resulted in a greater number of extended states. (A-C) In the corresponding three-state models, only *on, off*, and *in* states exist; that is, the *in* state is not futher specified as *hi* or *lo*.

**SI Table 1.** Calculation steps for constructing Petri nets from discrete time-series data and formalism for the naming places and transitions. (A) Petri nets are constructed from a time series dataset in which the discretized expression states for a given set of genes are listed for each cell and time point. The structure of this so-called Node Table is illustrated using the example of two cells, three genes, and seven time points. (B) Table of changes in expression states (state table), as used for the construction of Petri nets. This table is obtained by processing the time series dataset in (A) row by row. The first entry for each plasmodium in the state table corresponds to the entry in the Node Table for the first time point of the time series of the cell in consideration. Each time an expression state of one of the considered genes changes in a cell’s time series, a row is added to the state table. This row specifies the new state combination and the time at which the new state was detected. In other words, time points in which no change in states occured are not included in the state table. Each combination of the three genes’ expression states defines the name of the place in the Petri net that represents this combination of states. The order of the expression states by which a place is named corresponds to the order of the gene names by which the Petri net is named. Thus, the place named *lo_lo_hi* in the Petri net named *psgA_ligA_rgsA* represents a state in which *rgsA* is in the *hi* expression state while the other two genes are in the *lo* expression state. (C) Transits between states in individual cells and their representation as cell-specific transitions in the Petri net. Transitions between successive states in individual cells are determined from the state table in (B). The time at which a transit occured is defined as the time at which the new state was detected. Each transition that occurred in a single cell is represented as a corresponding transition in the Petri net. Depending on whether the Petri net is to be represented graphically using a Snoopy file or solely for computational analysis of its structure, we use two ways to name transitions. In the graphical representation, the name *t1_5*, for example, refers to the transition of cell number 1 that occurred five hours after the start of the experiment. For computational analysis, the same transition is designated *t1_5_lo_lo_hi_To_lo_on_off*. This means cell number one transitioned from the *lo_lo_hi* state to the *lo_on_off* state five hours after the start of the experiment. Since each transition name contains the pre- and post-place names, the complete set of transition names uniquely encodes the state machine structure represented by the Petri net. (D) Naming transitions in Petri nets in which parallel transitions are combined into a single transition. Rather than representing each transition of a single cell with a corresponding transition, the transits of all cells that occurred between two states in the same direction can be combined into a single transition. Note that this may not be formally correct from a biological perspective. The naming of these transitions lumping individual transits differs accordingly. In the graphical representation, the designation *t5_2x* means that transition number 5 occurred twice in the time series from which the Petri net was constructed. ID numbers (1 to n) uniquely identify transitions but do not contain additional information about individual cells or the time at which the transit occurred. In computational analyses, the name of each transition consists of the name of the pre-place and the name of the post-place of the transition. The complete set of transition names also uniquely defines the structure of the Petri net.

| (A) Time series of discrete Gene expression states (Node Table) |  |  |  |  |  |  |  |  |
| Cell | time *t* (h) | *State of each Gene* |  |  |  |  |  |  |
|  |  | *GeneA* | *GeneB* | *GeneC* |  |  |  |  |
|---|---|---|---|---|---|---|---|---|
| Cell 1 | 0 | lo | lo | hi |  |  |  |  |
| Cell 1 | 1 | lo | on | off |  |  |  |  |
| Cell 1 | 2 | lo | on | off |  |  |  |  |
| Cell 1 | 3 | lo | hi | off |  |  |  |  |
| Cell 1 | 4 | lo | hi | off |  |  |  |  |
| Cell 1 | 5 | on | hi | off |  |  |  |  |
| Cell 1 | 6 | on | hi | off |  |  |  |  |
| Cell 2 | 0 | lo | lo | hi |  |  |  |  |
| Cell 2 | 1 | lo | lo | hi |  |  |  |  |
| Cell 2 | 2 | lo | on | hi |  |  |  |  |
| Cell 2 | 3 | lo | on | hi |  |  |  |  |
| Cell 2 | 4 | lo | hi | off |  |  |  |  |
| Cell 2 | 5 | lo | hi | off |  |  |  |  |
| Cell 2 | 6 | on | hi | off |  |  |  |  |
| (B) Table of changes in expression states (State table) for PN construction |  |  |  |  |  |  |  |  |
| Cell | time *t* (h) | *State of each Gene* |  |  | *Name of* | *Tokens in places Gene* |  |  |
|  |  | *GeneA* | *GeneB* | *GeneC* | *PN Place* | *GeneA* | *GeneB* | *GeneC* |
| Cell 1 | 0 | lo | lo | hi | lo_lo_hi | 0 | 0 | 4 |
| Cell 1 | 1 | lo | on | off | lo_on_off | 0 | 3 | 1 |
| Cell 1 | 3 | lo | hi | off | lo_hi_off | 0 | 4 | 1 |
| Cell 1 | 5 | on | hi | off | on_hi_off | 3 | 4 | 1 |
| Cell 2 | 0 | lo | lo | hi | lo_lo_hi | 0 | 0 | 4 |
| Cell 2 | 2 | lo | on | hi | lo_on_hi | 0 | 3 | 4 |
| Cell 2 | 4 | lo | hi | off | lo_hi_off | 0 | 4 | 1 |
| Cell 2 | 6 | on | hi | off | on_hi_off | 3 | 4 | 1 |
| (C) Transits of states in individual cells represented as cell-specific Petri net transitions |  |  |  |  |  |  |  |  |
|  | Transit at | Names of places |  | Names of transitions |  |  |  |  |
| Cell | time *t* (h) | Pre-place | Post-place | in PN Graph | in computational representation |  |  |  |
| Cell 1 | 1 | lo_lo_hi | lo_on_off | t1_1 | t1_1_lo_lo_hi_To_lo_on_off |  |  |  |
| Cell 1 | 3 | lo_on_off | lo_hi_off | t1_3 | t1_3_lo_on_off_To_lo_hi_off |  |  |  |
| Cell 1 | 5 | lo_hi_off | on_hi_off | t1_5 | t1_5_lo_lo_hi_To_lo_on_off |  |  |  |
| Cell 2 | 2 | lo_lo_hi | lo_on_hi | t2_2 | t2_2_lo_lo_hi_To_lo_on_hi |  |  |  |
| Cell 2 | 4 | lo_on_hi | lo_hi_off | t2_4 | t2_4_lo_on_hi_To_lo_hi_off |  |  |  |
| Cell 2 | 6 | lo_hi_off | on_hi_off | t2_6 | t2_6_lo_hi_off_To_on_hi_off |  |  |  |
| (D) Individual transits of states lumped into common transitions in the Petri net |  |  |  |  |  |  |  |  |
| Names of places |  | Names of transitions |  |  |  |  |  |  |
| Pre-place | Post-place | in PN Graph* | in computational representation |  |  |  |  |  |
| lo_lo_hi | lo_on_off | t1_1x | lo_lo_hi_To_lo_on_off |  |  |  |  |  |
| lo_lo_hi | lo_on_hi | t2_1x | lo_lo_hi_To_lo_on_hi |  |  |  |  |  |
| lo_on_off | lo_hi_off | t3_1x | lo_on_off_To_lo_hi_off |  |  |  |  |  |
| lo_on_hi | lo_hi_off | t4_1x | lo_on_hi_To_lo_hi_off |  |  |  |  |  |
| lo_hi_off | on_hi_off | t5_2x | lo_lo_hi_To_lo_on_off |  |  |  |  |  |

\* name indicates how often a transit occurred (see legend for details)
Abbreviation: PN, Petri net

#### Random Gene Node Table

To evaluate the extent to which the analysis results are gene-specific, Random Gene Node Tables were generated from the Node Table. In these tables, the gene names of each trajectory are randomly assigned (SI Fig. 7). During randomization, the expression states and temporal patterns of changes remain unchanged for each trajectory so as to not artificially alter the number of state changes for a gene. Each trajectory of state changes occurring in a cell is assigned the name of a gene randomly and without replacement from all the analyzed gene names. Thus, each cell’s trajectories contain their own random gene name assignment. Setting a random seed makes the result of the randomization reproducible and allows for the generation and comparison of different randomized versions of a Random Gene Node Table. When shuffling the gene names, no distinction was made between up- and down-regulated genes since practically all genes exhibited all possible states or state changes.

#### Pseudo Cell Node Table

To examine the extent to which observed phenomena are cell-specific and thus vary individually, pseudo-cell trajectories were generated by randomly shuffling the plasmodium names at each time point in the time series of an individual cell. The expression values of all genes for a cell at each time point remained unchanged. The result is a pseudo-cell time series, where each time series consists of randomly selected gene expression patterns from a pool of cells without replacement while maintaining the original temporal order. Only irradiated plasmodia were used to shuffle the cell names (cell ID numbers) to generate the pseudo-cell trajectories. This ensures that no datasets are created in which gene expression time series from irradiated plasmodia are mixed with unexposed dark controls, as these groups exhibit different gene expression patterns. The shuffled cell names are applied in the same order to the mean and SEM values so each mean value is correctly assigned to its respective SEM value. This dataset serves as input for the double-arrow algorithm, which generates the Pseudo-Cell Node Table. The Extended State Pseudo-Cell

Node Table is derived from this table. SI Fig. 7 provides an overview of the computational steps and naming conventions for various types of Node Tables. The different types of Node Tables used in this study are provided as SI File 4.

**SI Figure 7.**
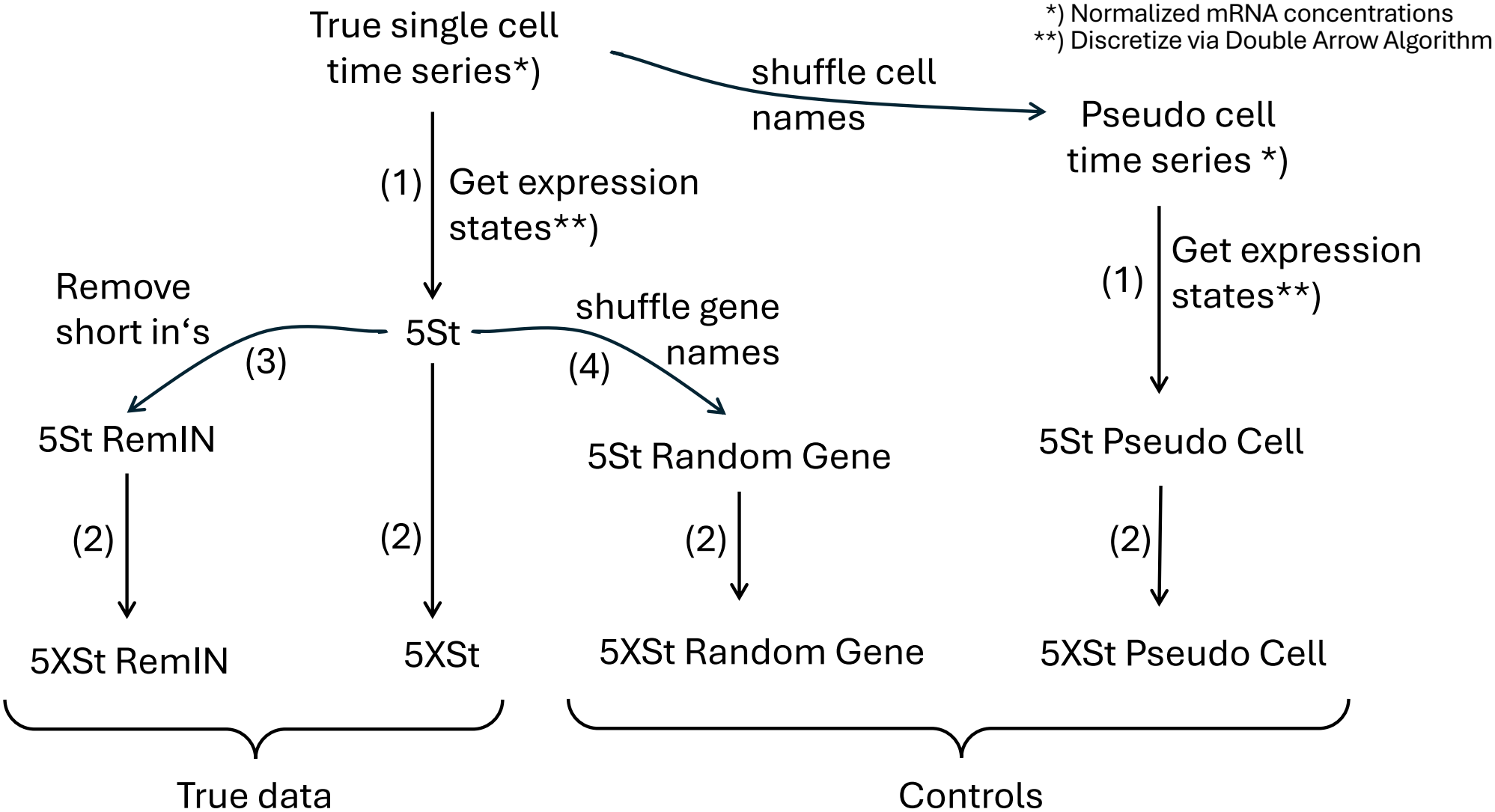
Generation of trajectories and pseudotrajectories and their annotation in corresponding types of Node Tables. The overview diagram illustrates the computational steps that begin with the longitudinal time series, which is the time course of relative mRNA concentrations in individual cells. The numbers in parentheses indicate the computational steps and point to the respective R code used. The scheme refers to the 5-state models but is valid for the family of 3-state models accordingly. First, the time course of gene expression states is determined from the time course of mRNA concentrations using the double-arrow algorithm. The trajectories, in the form of the temporal sequence of expression states in each cell, are stored in the five-state Node Table (5St). These trajectories can be analyzed or used as a basis for further processing. Identifying the extended states (2)—the states that persist over a longer period (i.e., a user-defined minimum time)—yields the 5XSt Node Table. Alternatively, short-lived *in* states can be removed from the 5St Node Table (3) if they occur between identical states, converting a short *in* state between two *on* states to *on* and a short *in* state between two *off* states to *off* (5StRemIN). Extended states can also be identified in the resulting trajectories (2), yielding a 5XStRemIN Node Table. To verify whether the identified effects are gene-specific, the gene names in a cell’s trajectory are randomly reassigned at each time point, preserving the temporal patterns of state changes while ensuring that each cell receives a distinct random assignment. This yields a Random Gene Node Table in which the extended states can be identified, resulting in the 5XSt random gene Node Table. To test the extent to which the observed effects arise from analyzing the distinct trajectories of individual cells, pseudo-cell time series are obtained from the time courses of relative mRNA concentrations in individual cells. This is done by randomly reassigning the original ID numbers of the cells for each time point of a time series (5). Thus, each pseudo-cell time series comprises data from randomly (without replacement) selected cells. The gene expression states in the pseudo cells are obtained using the double-arrow algorithm (1) according to the 5-state model (5St Pseudo Cell Node Table), and the 5XSt Cell Pseudo Node Table is generated by determining the extended states. The abbreviations characterizing the different types of Node Tables are used as name elements to specifiy the respective Node Table files in the computational pipeline (see SI Figure A2-1).

#### Construction of Petri Nets

Petri nets were constructed from a Node Table (see SI Table 1A) for a specific group of genes, called an x-plet, as illustrated in Figures 1 D-F. For graphical representation, the Petri nets were encoded in ANDL format (Abstract Net Description Language; (Heiner et al. 2013)) and exported as text. The program generates two versions of each Petri net: one in which each cell’s transitions are represented as individual, potentially parallel transitions and one in which all parallel transitions between two places are combined into a single transition. These transitions are named for graphical representation, as described in the legend for SI Table 1A. Each ANDL file was then imported into Snoopy (Heiner et al. 2012) to obtain the graphical layout of an executable stochastic Petri net. To enable the efficient batch processing of a large number of files, importing ANDL files into Snoopy, automatically laying out the files using the Sugiyama algorithm in Snoopy, and exporting the files in EPS format were automated by controlling Snoopy via a Python script accessed by an R script. To color-code places and transitions according to various criteria for visualization purposes, the text of the Snoopy XML files was edited automatically using an R script.

For computational analysis, each Petri net was encoded as a set of transition names in the format described in SI Table 1C,D. Each set of names was then evaluated for generating and structurally analyzing Petri nets.

The R code for the computational pipeline and the Python script for controlling Snoopy are provided as supplementary information.

#### Analysis of Hubs

Petri nets were used to identify hubs, where each transition of a cell from one gene expression state to another is represented by a cell-specific transition. By analyzing the transition names that describe the Petri net (SI Table 1C), all places with a minimum number of incoming edges were identified. For each place defined as a hub based on this minimum number, the following parameters were entered into a list for later evaluation: (1) the expression status of the genes represented by the hub; (2) the transitions leading to the hub; (3) the ID numbers of the cells in which the transition-generating transit occurred; (4) the time at which the transit took place; and (5) the number and identity of the pre-places directly connected to the hub. We then performed an analysis of the hubs and their properties by querying and evaluating this list (not shown).

### Supplementary Results

#### Temporal Order of Gene Expression

To identify genes entering an extended state, we first generated a Node Table of extended states from single-cell trajectories obtained from the time series using the double-arrow algorithm. In the Extended State Node Table, we included all states that persisted as extended states for at least three hours (SI Figs. 6 and 7). Then, the number of cells that entered an extended state over the course of the time series was determined for each gene from the trajectories obtained in this way. Based on this list, we selected genes for which at least 14 out of 16 cells entered an *Xon* state and at least 12 cells entered an *Xoff* state. This allowed us to obtain data for further analysis. We found that 14 out of the 72 differentially regulated genes analyzed entered an *Xon* state in at least 14 cells, and seven genes entered an *Xoff* state in at least 12 cells. Of these 21 genes, a significant temporal sequence was found for 13 *Xon* and six *Xoff* genes, respectively, relative to at least one other gene in this group (see below).

The time at which a gene first entered an extended state in a given cell was plotted in a Dumbbell plot. SI Fig. 8A shows when the various cells first entered an extended state on separate timelines for each gene. SI Fig. 8B shows the temporal pattern of extended state occurrence on separate timelines for each cell, allowing for an easy comparison of the behavior of different cells. Despite the relatively high variability in the temporal patterns, cascade-like regulation is clearly visible.

We investigated the extent to which the cascade-like pattern results from a strict temporal sequence in which genes initially enter an extended state (*Xon* or *Xoff*) at the earliest time point following stimulation. We performed a pairwise comparison to examine whether genes that entered an extended state in all or most cells did so in the same temporal order relative to one another or if the temporal order appeared random. The diagrams in SI Figs 9 and 10 show how the temporal order was algorithmically determined. For each gene pair, we evaluated the number of cells in which GeneA entered the extended state before GeneB and the number of cells in which GeneB entered the extended state before GeneA. If a gene did not reach an extended state during the observation period, we set its time of entry to infinity. Cases in which GeneA and GeneB entered simultaneously (ties) were noted and considered in significance tests based on sample size. We determined the significance of the temporal order for each gene pair using a binomial test and a permutation test. The permutation test does not rely on any model assumptions regarding the statistical frequency distribution of the data. The permutation test randomly swaps the names of the genes in a gene pair for each analyzed cell to examine the relative frequency with which the observed result is obtained purely by chance. We calculated the p-values of both tests using the Benjamini-Hochberg correction for multiple testing to determine the false discovery rate (FDR). Comparing the p-values obtained shows that both methods yield consistent results (SI Fig. 11).

**SI Figure 8.**
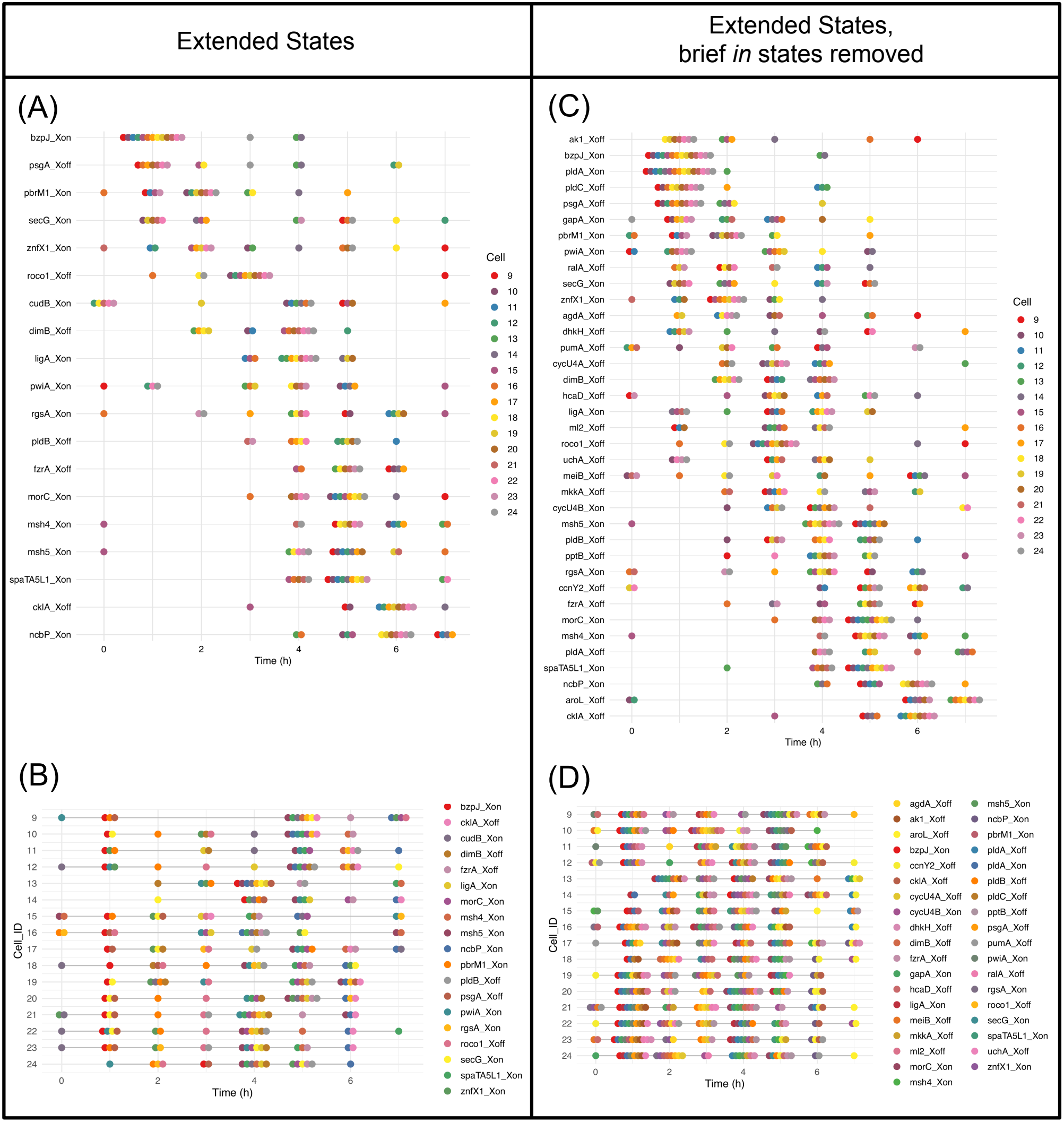
Dumbbell plots of the temporal patterns with which genes enter an extended state. Each data point shows the time at which an extended state initially occurred for each gene and cell in the time series. The figure compares the temporal patterns obtained by including (A, B) or removing (C, D) the short *in* states of one hour length at maximum. Panels A and C show, on separate timelines for each gene, when the various cells first entered an extended state. Panels B and D show the temporal pattern of extended state occurrence on separate timelines for each cell to compare the behavior of different cells.

**SI Figure 9.**
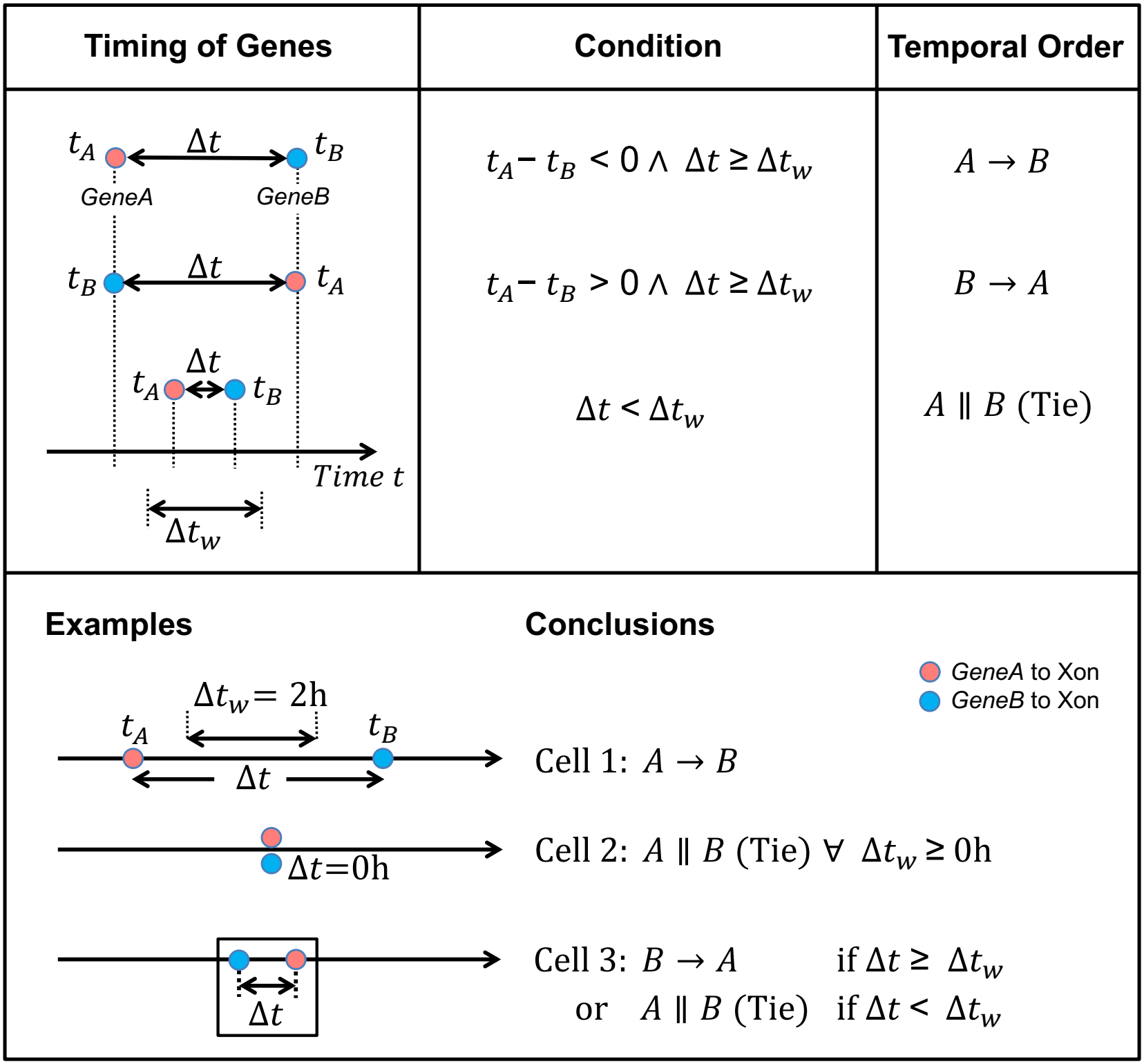
Determining the temporal order in which two genes, GeneA and GeneB, entered an extended state. For brevity, entering an extended state is abbreviated as “regulated” in the context of this figure legend. The diagram shows how the algorithm compares time points *t*_*A*_ and *t*_*B*_, at which the respective gene entered the extended state. The algorithm distinguishes between three cases: 1) GeneA is regulated before GeneB (A->B), 2) GeneB is regulated before GeneA (B->A), or 3) both genes appear within the same time window (i.e., they are regulated relatively simultaneously, A||B). The width of the user-defined time window (*Δt*_*w*_) determines the minimum time interval between the entry of two genes into an extended state that must be observed for a temporal sequence to be recognized. In the analyzed time series with hourly sampling, *Δt*_*w*_ values of 1, 2, or 3 mean that the genes under consideration had to enter the extended state with relative delay times of at least 1, 2, or 3 hours to be classified as sequentially regulated. Thus, the variation in the width of the time window serves sensitivity analysis and reflects the magnitude of the time difference in differential regulation. The lower part of the figure shows three cells in which Gene A and Gene B enter the extended state with different temporal patterns. In Cell 1, the time interval is greater than *Δt*_*w*_, so the expression state of GeneA is assumed to change before that of GeneB. In Cell 2, the regulation (entry into an extended state) occurs simultaneously, independent of *Δt*_*w*_. For Cell 3, however, the interpretation of the result depends on the width of the time window, *Δt*_*w*_. The value of *Δt*_*w*_ determines whether genes A and B in Cell 3 appear to be expressed simultaneously or in reverse order as compared to Cell 1.

**SI Figure 10.**
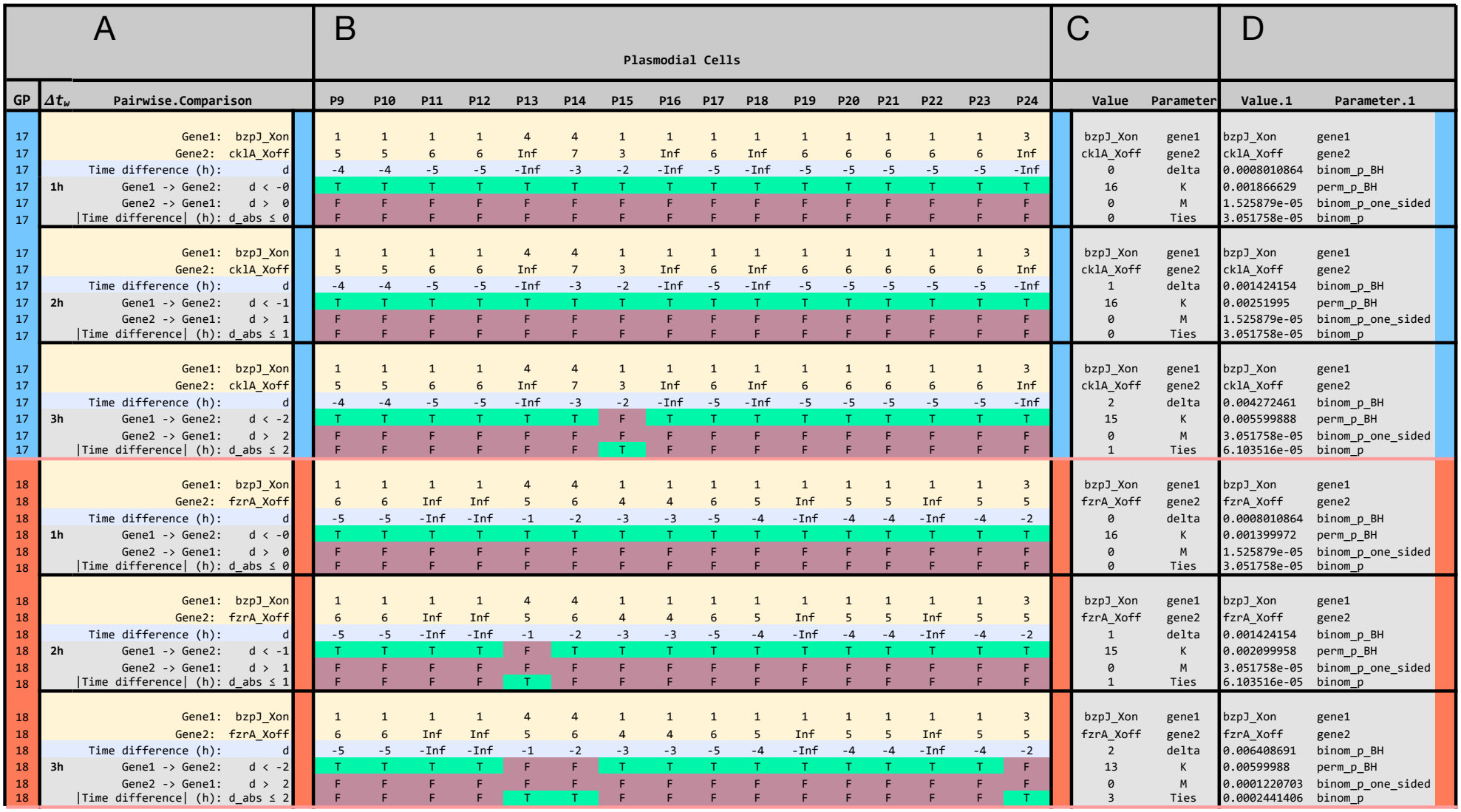
Analyses and intermediate results of the test for temporal order of entry into an extended state. The diagram illustrates how the test determines if two genes enter an extended state in a significant temporal order, showing two gene pairs as example. The data of each gene pair are shown in three consecutive horizontal blocks (*Δt*_*w*_ = 1h, 2h, 3h), each block showing the results obtained for the given gene pair across the time window *Δt*_*w*_. Let us consider the first horizontal block to explain terminology and abbreviations. The first two rows in vertical block B, *Plasmodial Cells*, show the earliest times, in hours, after stimulation at which each of the two genes, *bzpJ* and *cklA*, entered an extended state in each cell (P9 through P24). In cases where no extended state occurred, this time is set to infinity (Inf). The third row specifies the time difference *d* between Gene1 and Gene2 entering the extended state. The true/false values in lines four to six indicate whether Gene1 is regulated before Gene2 or *vice versa*, or whether the two genes are not regulated in a temporal order relative to each other. In other words, a temporal order is assumed if the time difference (d) between the first entry of two genes into an extended state after stimulation is at least *Δt*_*w*_ (see also SI Fig. 9). As shown in the example for cell P15, the algorithm only identifies *bzpJ* as entering an extended state before *cklA* when the time window *Δt*_*w*_ set as the criterion is two hours or less. Otherwise, the two genes are considered ties. Simultaneous changes in expression state are indicated by the true/false values in row 6. The values of the corresponing parameters in vertical block C indicate the width of the time window (expressed in *d* with *d* = *Δt*_*w*_ - 1h), for which the evaluation was performed, as well as the number of plasmodial cells in which Gene1 (M) entered the extended state before Gene2 (K), Gene2 entered the extended state before Gene1, or both genes entered the extended state simultaneously (Ties). Block D specifies the p-values of the two-sided binomial and permutation tests, with and without Benjamini-Hochberg correction, as well as the p-values of the one-sided and two-sided binomial test. The redundant gene and state names confirm the consistent assignment of the intermediate results of the calculations, which were automatically compiled into the shown table. A complete, searchable list of the results for all tested gene pairs displayed in this format is included in SI File 5.

Sensitivity analyses were conducted regarding the time window that defines simultaneous entry of both genes into the extended state. Evaluations were performed for time windows of 1, 2, and 3 hours (SI Figs. 9,10,12). The evaluation scheme and the results obtained for the different time windows in each cell are shown in SI Fig. 10, illustrated with genes that are regulated in a defined temporal order. The entire dataset is presented in this form in SI File 5.

**SI Figure 11.**
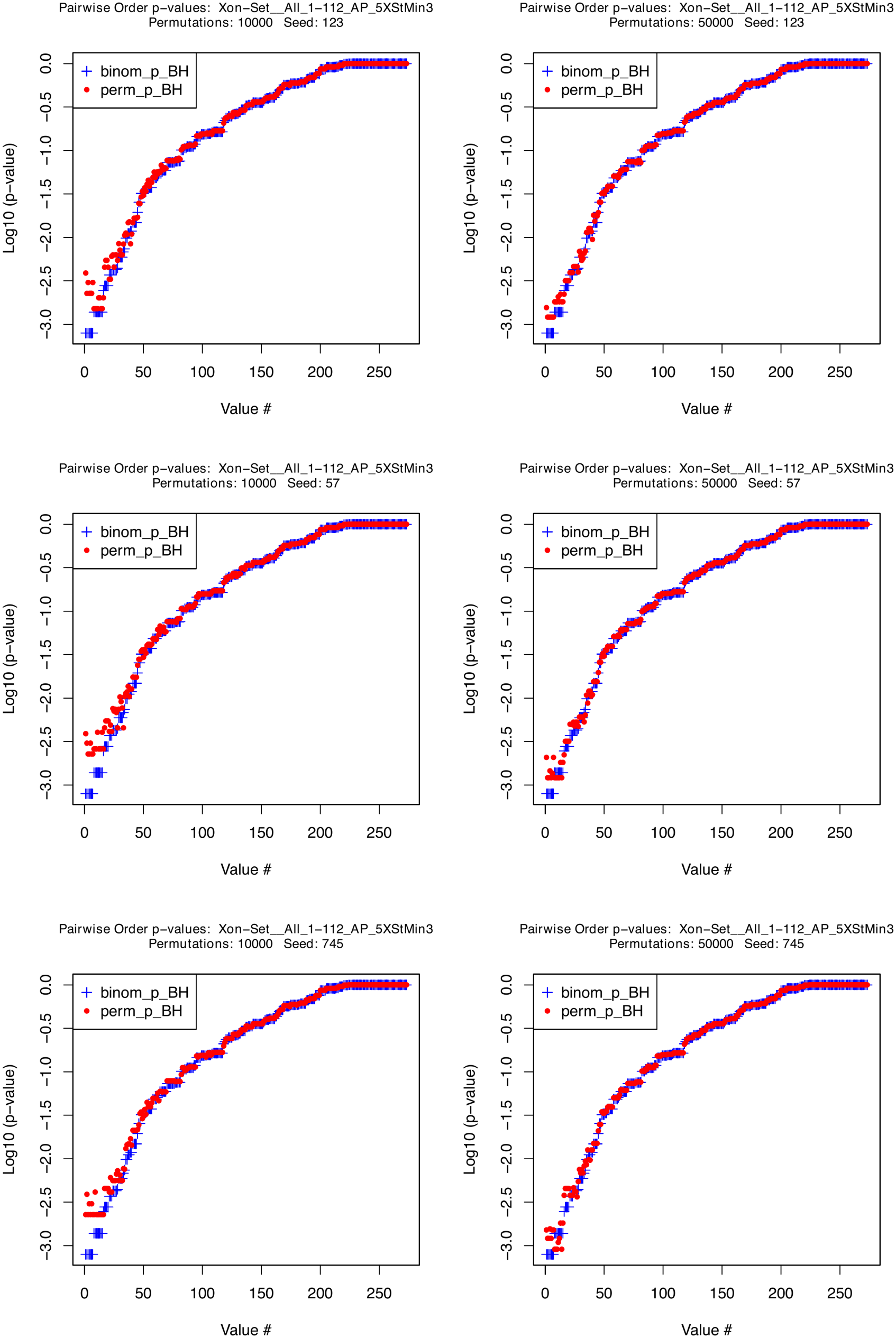
Comparison of p-values from the binomial test and the permutation test. This figure displays the p-values adjusted using the Benjamini-Hochberg correction for the binomial and permutation tests of all analyzed gene combinations in the *Xon* set. The permutation tests shown in the various panels were performed with different numbers of permutations and random seeds, as indicated in the panel headings.

**SI Figure 12.**
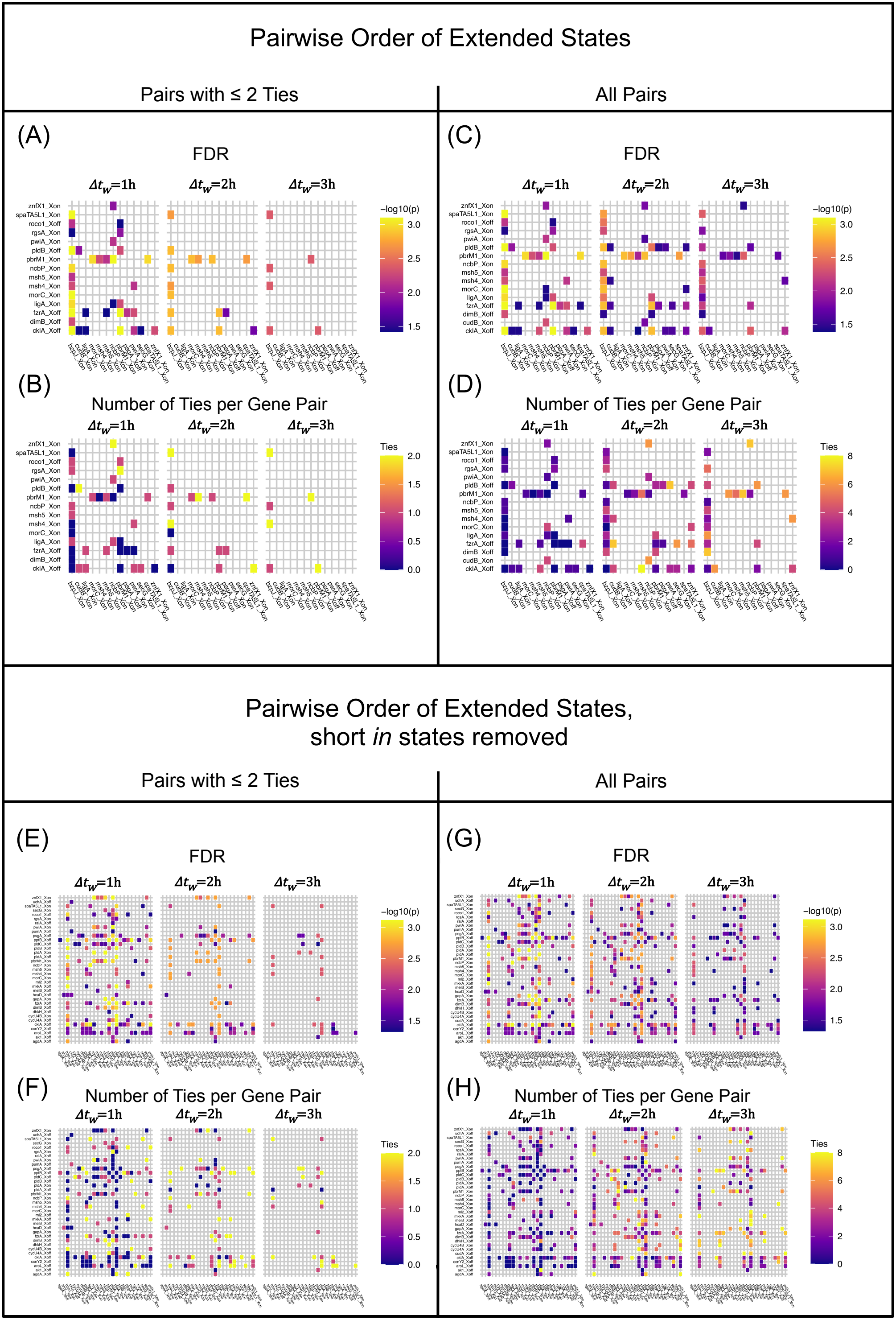
Gene pairs that entered an extended state in a significant temporal order. The figure shows combinations of two heatmaps, each representing the results obtained for different parameter settings in the analysis of the dataset, as indicated. The upper heatmaps (A, C, E, G) show the p-values of the binomial test for the significant temporal order of each gene pair, corrected for multiple testing using the Benjamini–Hochberg method, with a false discovery rate (FDR) of ≤0.05. The lower heatmaps (B, D, F, H) of each combination show the absolute number of ties found for each gene pair, equivalent to the number of cells in which the two genes entered the extended state simultaneously.

**SI Figure 13.**
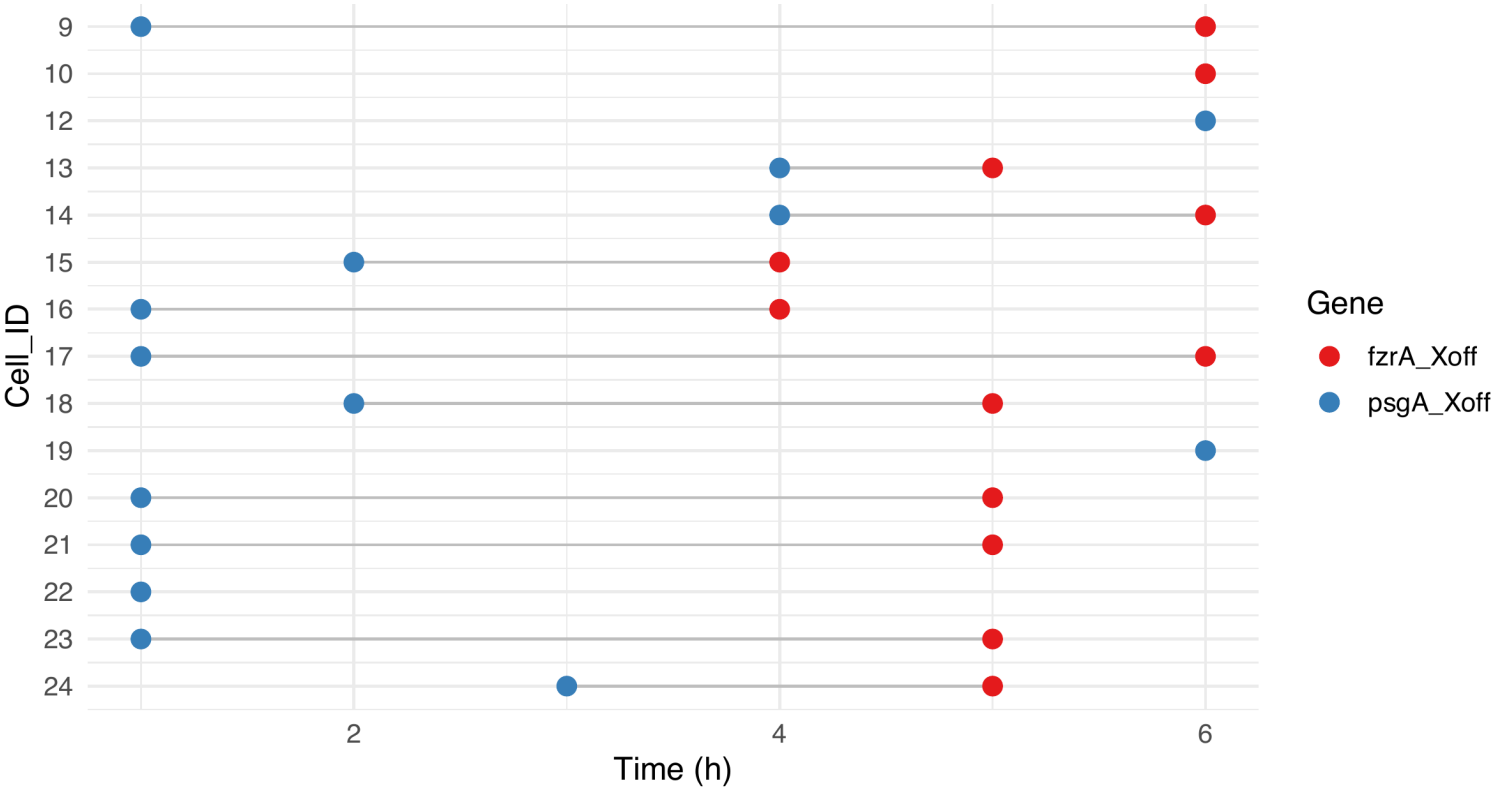
Individual differences in the timing of entering an extended state despite the same temporal order of the two genes. The figure shows a Dumbbell plot, where each data point represents the time of entering an extended state, with a separate timeline for each cell.

The p-values obtained using the binomial test with Benjamini-Hochberg correction for the temporal order in which two genes enter an extended state following stimulation are shown in the heatmaps in SI Fig. 12 A,C. Many pairs are evident in which Gene A is regulated in a significant temporal order relative to Gene B. The heatmaps identify genes that entered an extended state in a significant temporal order relative to a relatively large number of other genes based on the tiles lying on the same vertical or horizontal line.

Comparing the heatmap of p-values with the heatmap showing the number of ties per gene pair (SI Fig. 12 B,D) reveals that some gene pairs have a significant temporal order of entry into an extended state, even if one or more ties occurred within the pair. The presence of a significant temporal order and the occurrence of ties between the two genes suggests that Gene A enters an extended state before Gene B, or at most simultaneously with Gene B, but not vice versa. An extended state of GeneA could be required for the entry of GeneB into an extended state, but the reverse is not true. The same holds true if hypothetical regulators X and Y are activated in a causal sequence (X → Y) and GeneA depends on X and GeneB depends on Y; in this case, GeneA cannot enter an extended state before GeneB, at most entering within the same time window.

The variability of gene expression despite a defined temporal sequence is clearly demonstrated by the example of *psgA* and *fzrA*. Despite the strict temporal sequence in which both genes enter the *Xoff* state in all analyzed cells, the chronological times relative to the application of the triggering stimulus vary greatly from cell to cell (SI Fig. 13). A strict temporal sequence is also observed for other gene pairs, with high variability regarding the chronological time points from cell to cell.

**SI Figure 14.**
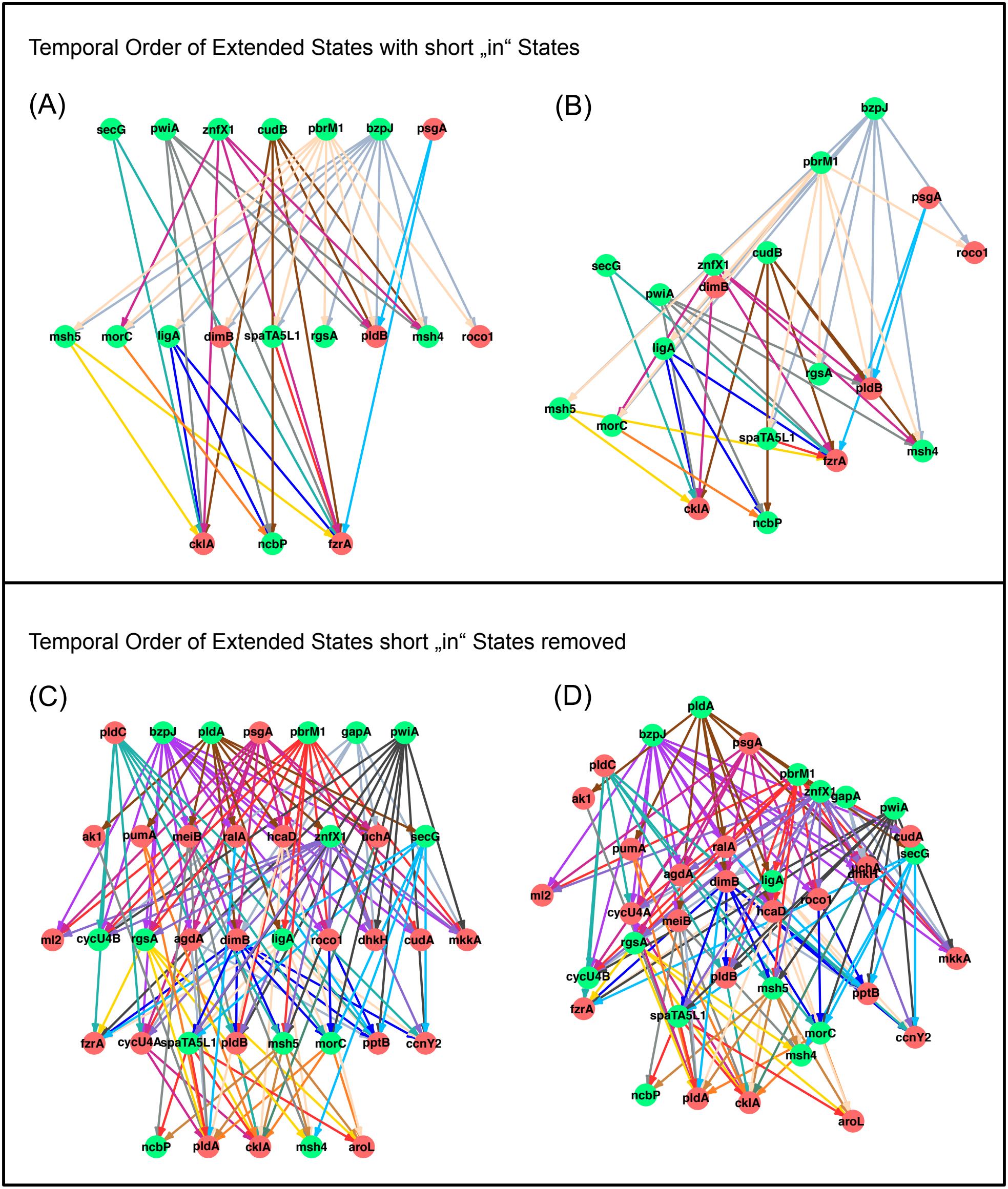
The temporal order in which genes enter an extended state, depicted as iGraph plots using the Sugiyama layout. The directed arcs between the nodes illustrate the temporal order in which two genes transitioned to an *Xon* (green) or *Xoff* (red) state. The iGraph plots depict the temporal order of gene pairs without (A, B) or after removing the short *in* states (C, D). Each graph is shown in two different layouts. The graphs in (A, C) are displayed in plain Sugiyama layout. The time-coded graphs in (B, D) were generated by modifying the original Sugiyama layout so that the vertical arrangement of the nodes corresponds to the mean of the delay times after stimulation at which the respective gene entered an extended state. The node representing the earliest gene is placed at the top, and the node representing the latest gene is placed at the bottom.

#### Temporal order of gene regulation displayed in Sugiyama layout

From the gene pairs that entered an extended state in a statistically significant temporal order, a single connected graph was obtained using the iGraph package (SI Fig. 14). The directed arcs connecting two nodes represent the temporal order in which the genes entered the extended state. The graph was checked for cycles, which would indicate contradictions in the temporal order of the depicted genes. For example, a cycle would arise if GeneA were activated before GeneB and GeneB before GeneA. Redundant edges with respect to the temporal order were removed. If GeneA entered an extended state before GeneB, and then GeneB entered an extended state before GeneC, for example, the graph would contain the edges A->B and B->C, but not the redundant edge A->C.

After removing the redundant edges, the graph was plotted using the Sugiyama layout to visualize the stepwise development of the system. The Sugiyama layout represents the hierarchical organization of the nodes, which are clearly arranged in successive, equidistant horizontal planes (SI Fig. 14 A,C). To roughly approximate the temporal development of the system, the y-coordinates of the original Sugiyama layout were modified so that the positions of the nodes on the y-axis correspond to the mean time after stimulus application at which the respective gene entered the extended state. In this arrangement, the earliest gene is placed at the top and the latest gene is placed at the bottom (SI Fig. 14 B,D). To determine the number of genes that entered an extended state consecutively in a significant temporal sequence relative to the subsequent gene, all longest unbranched paths through the graph were identified. For example, if a path branches once within a subgraph, two unbranched paths are obtained from this path. These unbranched paths are then sorted in descending order by the number of intermediates and output together with statistical information. SI Table 2 shows the longest unbranched paths through the graph. SI Table 3 shows the frequency in which genes and their extended states occur in the paths, quantifying the varying connectivity between individual genes. Complete versions of the tables from SI Tables 2 and 3 were generated can be found in SI Files 6,7.

#### Removing brief in states increases the apparent number of extended states and the number of pairwise-ordered genes

Based on considerations in the main manuscript about the unclear physiological relevance of short-lived *in* states during a time series, *in* states lasting no more than one hour that interrupted phases of decline or increase in mRNA concentration were removed from the Node Table. This included sequences of *on* -> *in* -> *on* to *on* and *off* -> *in* -> *off* to *off* (SI Fig. 6). All other *in* states were retained in the Node Table. Using the edited Node Table, we repeated the search for extended states and the analysis of the temporal order of entry into extended states. Removing the short *in* states increased the number of genes with an extended state from 19 to 36 for *Δt*_*w*_= 0 h.

Gene *pldA* assumed both an *Xon* and a later-occurring *Xoff* state (SI Fig. 8 C,D). The number of gene pairs for which a significant temporal order of entry into an extended state was found increased from 41 to 205 (FDR ≤ 0.05; SI Fig. 12 E–H). Moreover, the maximum number of genes entering an extended state sequentially in a significant temporal order increased from three to five when the short *in* states were removed, corresponding to the longest unbranched paths through the iGraph plot (SI Fig. 14 C,D, SI Files 6,7).

#### Random shuffling of gene names: Random-Gene Node Table

To test whether an established temporal order is gene-specific, we performed an additional control using a Random Gene Node Table, an approach separate from binomial and permutation tests. This table was obtained by randomly shuffling the gene names at each time point in the time series while maintaining the temporal patterns of gene expression and cell identity (see section on Node Tables for details). We searched for extended states in these randomized data and repeated the pairwise search for a temporal sequence of entry into an extended state. During the search for extended states, no genes were found in which nearly all or all cells exhibited extended states. Instead, extended states occurred in no more than 8–9 out of 16 cells, suggesting that this small number is a consequence of randomly reassigning gene names. SI Table 4 presents the number of gene pairs that were detected to enter extended states in a significant temporal order according to binomial and permutation tests, comparing various Node Tables and various p-values for real and pseudo-trajectories. With the Random Gene and Pseudo Cell Node Tables, no gene pairs with significant temporal order were found.

**SI Table 2.**
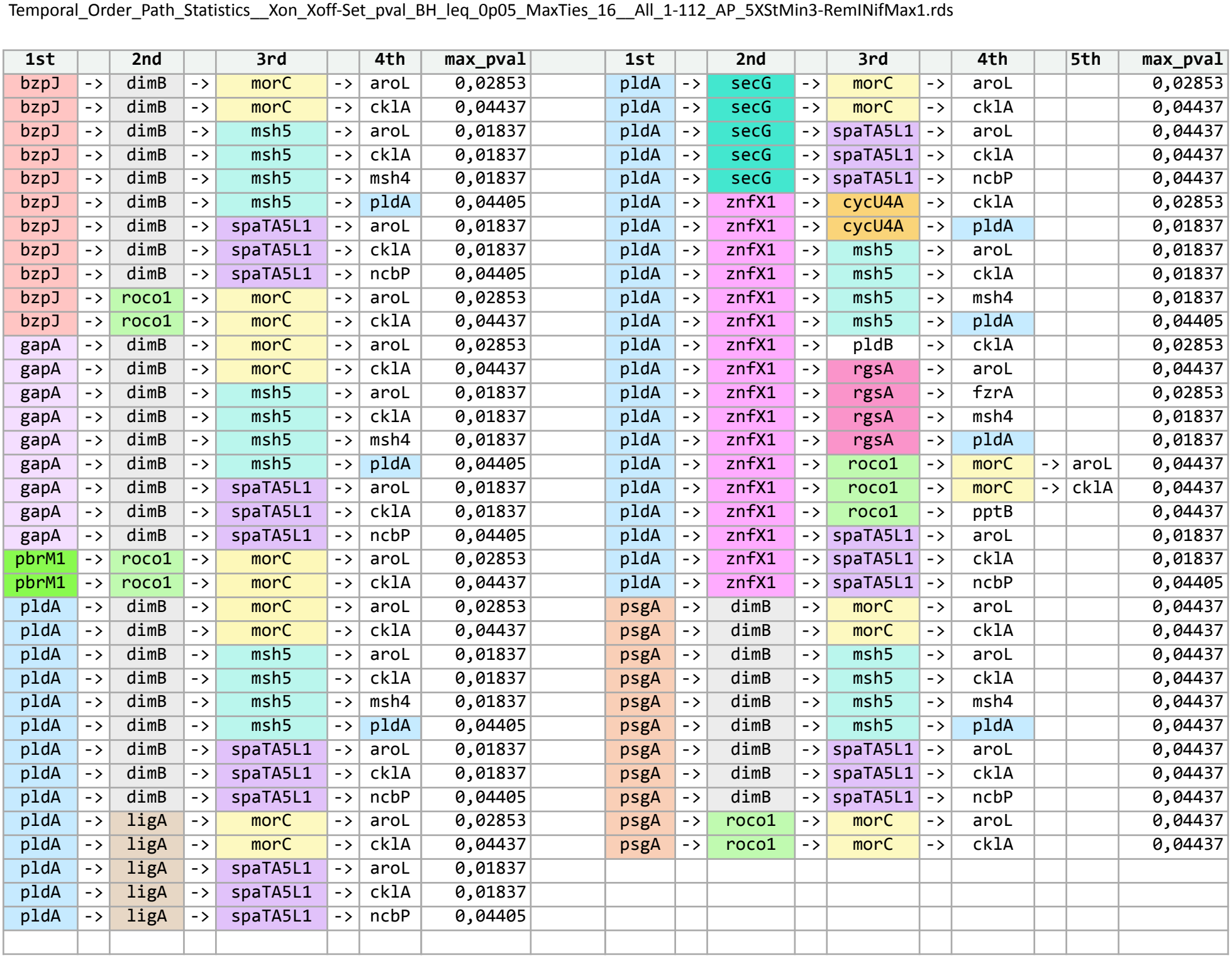
Genes in temporal order. The table shows the statistically significant temporal order (FDR “0.05 for each step) in which genes entered an extended state, as determined by the longest single paths through the graph in SI Fig. 14C. The Table shows only those paths that consist of at least four genes.

| 1st |  | 2nd |  | 3rd |  | 4th | max_pval |  | 1st |  | 2nd |  | 3rd |  | 4th |  | 5th | max_pval |
|---|---|---|---|---|---|---|---|---|---|---|---|---|---|---|---|---|---|---|
| bzpJ | -> | dimB | -> | morC | -> | aroL | 0,02853 |  | pldA | -> | secG | -> | morC | -> | aroL |  |  | 0,02853 |
| bzpJ | -> | dimB | -> | morC | -> | cklA | 0,04437 |  | pldA | -> | secG | -> | morC | -> | cklA |  |  | 0,04437 |
| bzpJ | -> | dimB | -> | msh5 | -> | aroL | 0,01837 |  | pldA | -> | secG | -> | spaTA5L1 | -> | aroL |  |  | 0,04437 |
| bzpJ | -> | dimB | -> | msh5 | -> | cklA | 0,01837 |  | pldA | -> | secG | -> | spaTA5L1 | -> | cklA |  |  | 0,04437 |
| bzpJ | -> | dimB | -> | msh5 | -> | msh4 | 0,01837 |  | pldA | -> | secG | -> | spaTA5L1 | -> | ncbP |  |  | 0,04437 |
| bzpJ | -> | dimB | -> | msh5 | -> | pldA | 0,04405 |  | pldA | -> | znfX1 | -> | cycU4A | -> | cklA |  |  | 0,02853 |
| bzpJ | -> | dimB | -> | spaTA5L1 | -> | aroL | 0,01837 |  | pldA | -> | znfX1 | -> | cycU4A | -> | pldA |  |  | 0,01837 |
| bzpJ | -> | dimB | -> | spaTA5L1 | -> | cklA | 0,01837 |  | pldA | -> | znfX1 | -> | msh5 | -> | aroL |  |  | 0,01837 |
| bzpJ | -> | dimB | -> | spaTA5L1 | -> | ncbP | 0,04405 |  | pldA | -> | znfX1 | -> | msh5 | -> | cklA |  |  | 0,01837 |
| bzpJ | -> | roco1 | -> | morC | -> | aroL | 0,02853 |  | pldA | -> | znfX1 | -> | msh5 | -> | msh4 |  |  | 0,01837 |
| bzpJ | -> | roco1 | -> | morC | -> | cklA | 0,04437 |  | pldA | -> | znfX1 | -> | msh5 | -> | pldA |  |  | 0,04405 |
| gapA | -> | dimB | -> | morC | -> | aroL | 0,02853 |  | pldA | -> | znfX1 | -> | pldB | -> | cklA |  |  | 0,02853 |
| gapA | -> | dimB | -> | morC | -> | cklA | 0,04437 |  | pldA | -> | znfX1 | -> | rgsA | -> | aroL |  |  | 0,04437 |
| gapA | -> | dimB | -> | msh5 | -> | aroL | 0,01837 |  | pldA | -> | znfX1 | -> | rgsA | -> | fzrA |  |  | 0,02853 |
| gapA | -> | dimB | -> | msh5 | -> | cklA | 0,01837 |  | pldA | -> | znfX1 | -> | rgsA | -> | msh4 |  |  | 0,01837 |
| gapA | -> | dimB | -> | msh5 | -> | msh4 | 0,01837 |  | pldA | -> | znfX1 | -> | rgsA | -> | pldA |  |  | 0,01837 |
| gapA | -> | dimB | -> | msh5 | -> | pldA | 0,04405 |  | pldA | -> | znfX1 | -> | roco1 | -> | morC | -> | aroL | 0,04437 |
| gapA | -> | dimB | -> | spaTA5L1 | -> | aroL | 0,01837 |  | pldA | -> | znfX1 | -> | roco1 | -> | morC | -> | cklA | 0,04437 |
| gapA | -> | dimB | -> | spaTA5L1 | -> | cklA | 0,01837 |  | pldA | -> | znfX1 | -> | roco1 | -> | pptB |  |  | 0,04437 |
| gapA | -> | dimB | -> | spaTA5L1 | -> | ncbP | 0,04405 |  | pldA | -> | znfX1 | -> | spaTA5L1 | -> | aroL |  |  | 0,01837 |
| pbrM1 | -> | roco1 | -> | morC | -> | aroL | 0,02853 |  | pldA | -> | znfX1 | -> | spaTA5L1 | -> | cklA |  |  | 0,01837 |
| pbrM1 | -> | roco1 | -> | morC | -> | cklA | 0,04437 |  | pldA | -> | znfX1 | -> | spaTA5L1 | -> | ncbP |  |  | 0,04405 |
| pldA | -> | dimB | -> | morC | -> | aroL | 0,02853 |  | psgA | -> | dimB | -> | morC | -> | aroL |  |  | 0,04437 |
| pldA | -> | dimB | -> | morC | -> | cklA | 0,04437 |  | psgA | -> | dimB | -> | morC | -> | cklA |  |  | 0,04437 |
| pldA | -> | dimB | -> | msh5 | -> | aroL | 0,01837 |  | psgA | -> | dimB | -> | msh5 | -> | aroL |  |  | 0,04437 |
| pldA | -> | dimB | -> | msh5 | -> | cklA | 0,01837 |  | psgA | -> | dimB | -> | msh5 | -> | cklA |  |  | 0,04437 |
| pldA | -> | dimB | -> | msh5 | -> | msh4 | 0,01837 |  | psgA | -> | dimB | -> | msh5 | -> | msh4 |  |  | 0,04437 |
| pldA | -> | dimB | -> | msh5 | -> | pldA | 0,04405 |  | psgA | -> | dimB | -> | msh5 | -> | pldA |  |  | 0,04437 |
| pldA | -> | dimB | -> | spaTA5L1 | -> | aroL | 0,01837 |  | psgA | -> | dimB | -> | spaTA5L1 | -> | aroL |  |  | 0,04437 |
| pldA | -> | dimB | -> | spaTA5L1 | -> | cklA | 0,01837 |  | psgA | -> | dimB | -> | spaTA5L1 | -> | cklA |  |  | 0,04437 |
| pldA | -> | dimB | -> | spaTA5L1 | -> | ncbP | 0,04405 |  | psgA | -> | dimB | -> | spaTA5L1 | -> | ncbP |  |  | 0,04437 |
| pldA | -> | ligA | -> | morC | -> | aroL | 0,02853 |  | psgA | -> | roco1 | -> | morC | -> | aroL |  |  | 0,04437 |
| pldA | -> | ligA | -> | morC | -> | cklA | 0,04437 |  | psgA | -> | roco1 | -> | morC | -> | cklA |  |  | 0,04437 |
| pldA | -> | ligA | -> | spaTA5L1 | -> | aroL | 0,01837 |  |  |  |  |  |  |  |  |  |  |  |
| pldA | -> | ligA | -> | spaTA5L1 | -> | cklA | 0,01837 |  |  |  |  |  |  |  |  |  |  |  |
| pldA | -> | ligA | -> | spaTA5L1 | -> | ncbP | 0,04405 |  |  |  |  |  |  |  |  |  |  |  |

#### Pseudo-Cell Time Series

When analyzing the longitudinal time series of individual cells in our dataset, we found a relatively strict temporal order of entry into an extended state for some genes. However, we also found marked variability in the chronological timing that occurred from cell to cell. Thus, detecting a temporal order may require examining individual cells. To test this hypothesis, we generated pseudo-cell time series from true time series data. In these pseudo-cell time series, the ID numbers of each cell were randomly and without replacement assigned to each time point in the time series, while the gene expression patterns in the form of mRNA concentrations at each time point and the time points themselves remained unchanged (see the Computational Methods section for details). Each pseudo-cell time series is accordingly composed of data from different cells. We searched for extended states in the resulting pseudo-cell time series and performed the aforementioned analysis of the temporal order of entry into an extended state. As a result, virtually no genes appeared to be expressed in a significant temporal order. The results obtained for the real and pseudo-cell time series are compared in SI Table 4.

**SI Table 3.**
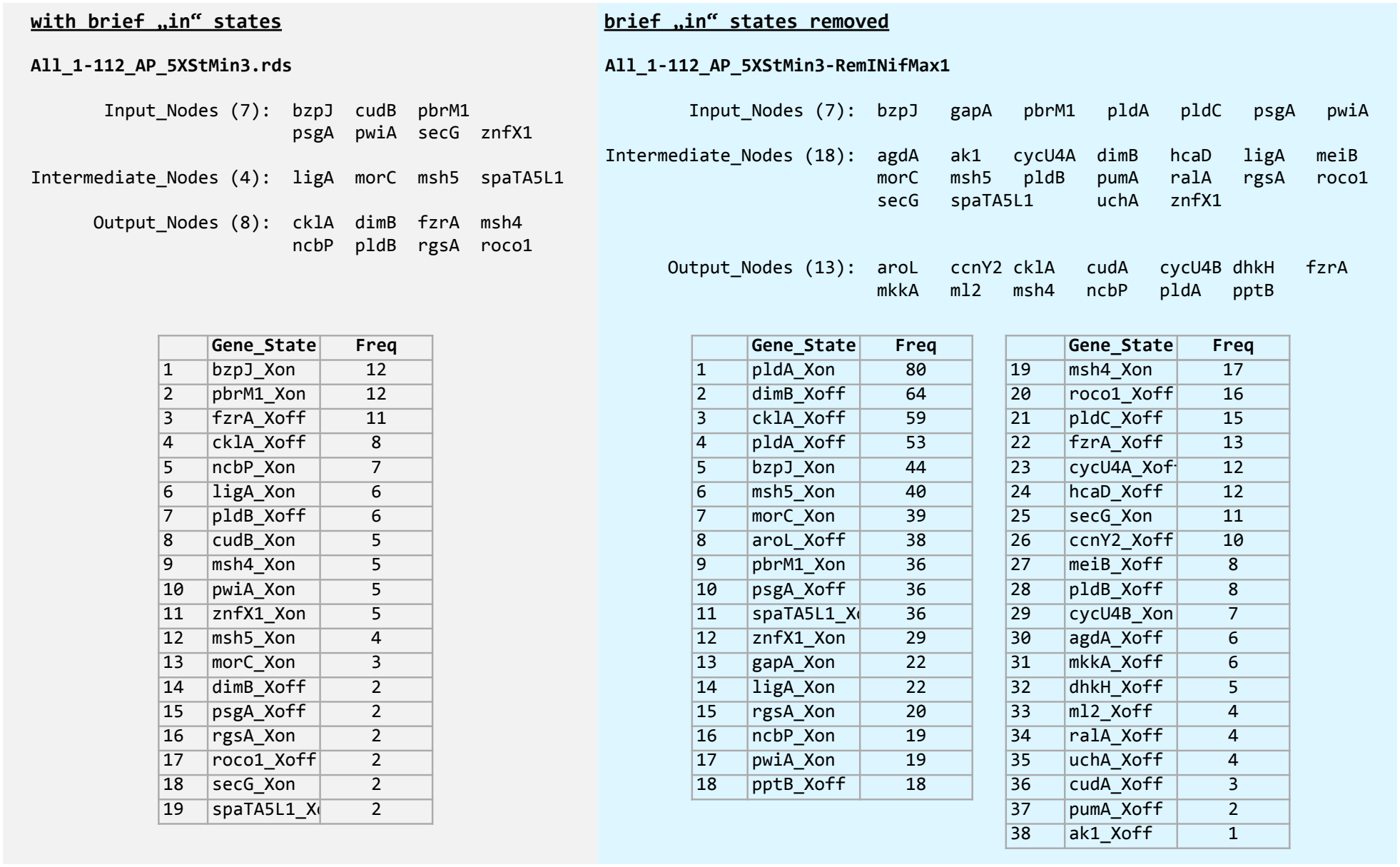
Absolute frequencies in which the extended states of genes occurred in paths through the Sugiyama plots of SI Figure 14, quantifying the varying connectivity of individual genes within the plot.

**SI Table 4.** Temporal order of genes entering an extended state found in trajectories of Random Gene and Pseudo Cell Node Tables as compared to those found for true cell trajectories. All possible combinations of *Xon* and *Xoff* states were considered. Node Tables used are provided in SI File 4.

| Node-Table | True data & controls | States considered | Number of gene pairs with significant temporal order (binomial p) |  |  |  |
|---|---|---|---|---|---|---|
|  |  |  | p ≤ 0.01 | p ≤ 0.025 | p ≤ 0.01<br>(one sided) | p ≤ 0.025<br>(one sided) |
| All_1-112_AP_5XStMin3.xlsx | True data | Xon, Xoff | 31 | 47 | 39 | 56 |
| All_1-112_AP_5XStMin3-RemINifMax1 | True data | Xon, Xoff | 119 | 148 | 133 | 178 |
| All_1-112_AP_5XStMin3_rs117_Random_Gene | Gene names shuffeled | Xon, Xoff | 0 | 0 | 0 | 0 |
| All_1-112_AP_5XStMin3-RemINifMax1_rs117_Random_Gene |  | Xon, Xoff | 0 | 0 | 0 | 0 |
| All_1-112_AP_5XStMin3-RemINifMax1_rs5_Random_Gene |  | Xon, Xoff | 0 | 0 | 0 | 2 |
| All_1-112_AP_5XStMin3_Pseudo_Cell_Node_Table.xlsx | Cell IDs shuffeled | Xon, Xoff | 0 | 0 | 0 | 0 |

**SI Figure 15.**
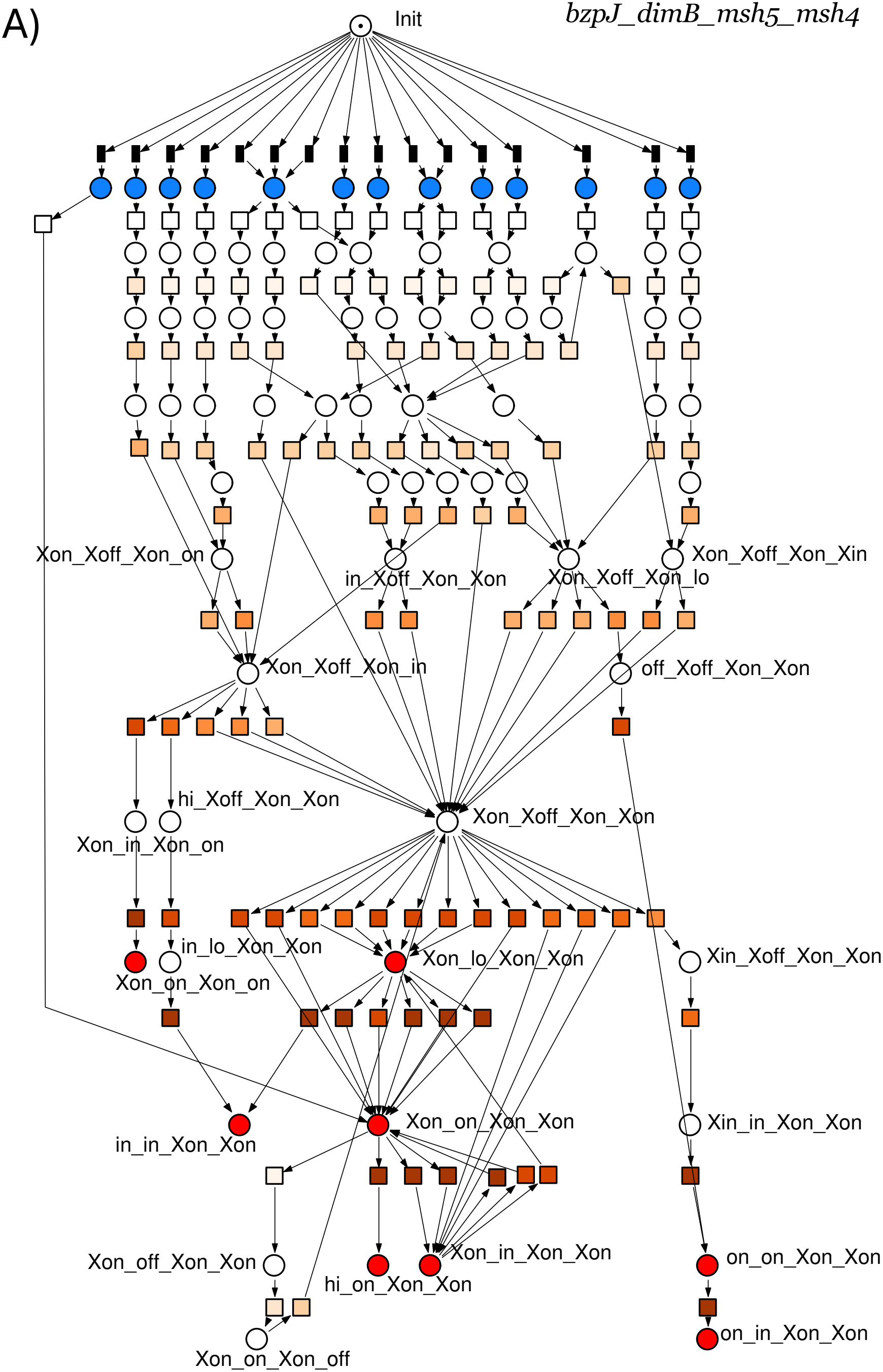
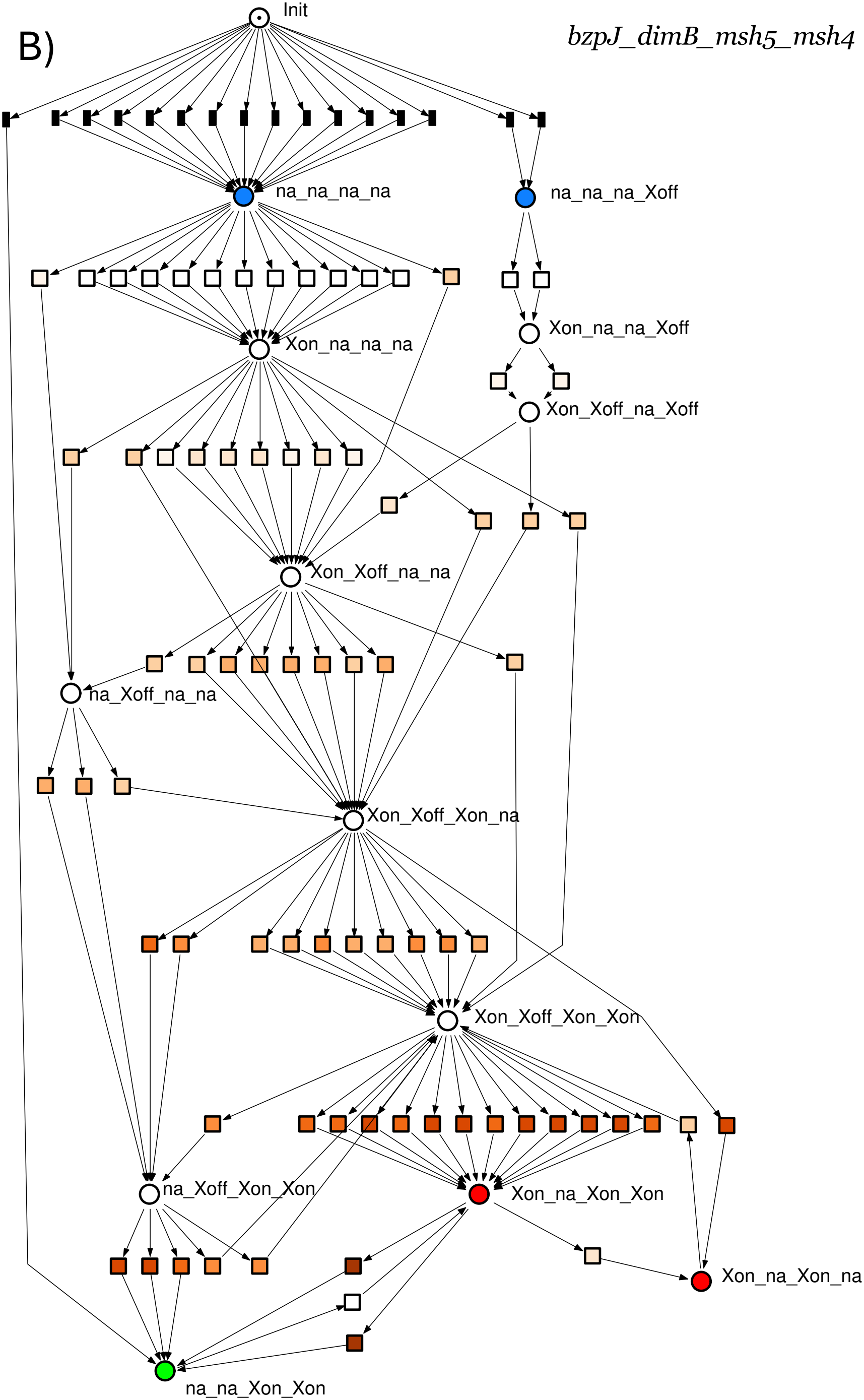
Petri net representation of the temporal order in which genes entered an extended state. Petri nets were constructed for the genes *bzpJ, dimB, msh5*, and *msh4*, which entered the extended state in a temporal order corresponding to an unbranched path through the iGraph plot (see SI Fig. 14C and 14D). The Petri nets were constructed using an extended-state Node Table obtained by removing short *in* states, as described. In the Petri net shown in (A), each place represents a combination of the observed expression states of the four genes. In the Petri net shown in (B), all states that are neither *Xon* nor *Xoff* are lumped into a single “na” state. The color coding of the transitions corresponds to the time of transit between two states, with the color intensity increasing with the time at which a switching process occurred.

**SI Table 5.** Functional annotation of genes that enter extended states based on highest sequence similarties to proteins in the UniProt database.

|  | **Transcript** | **Gene** | **Definition** | **Organism** | **UniProt** | **E.Value** | **Identity (%)** | **Length** |
|---|---|---|---|---|---|---|---|---|
| 1 | Phypoly_transcript_10011 | *ak1* | Alpha-protein kinase 1 | *Dictyostelium discoideum* | Q54DK4 | 2E-12 | 32 | 1352 |
| 2 | Phypoly_transcript_07191 | *fzrA* | Anaphase-promoting complex subunit cdc20 | *Dictyostelium discoideum* | Q54MZ3 | 1E-140 | 50 | 499 |
| 3 | Phypoly_transcript_03573 | *spaTA5L1* | ATPase family gene 2 protein homolog B | *Rattus norvegicus* | D4A2B7 | 2E-137 | 40 | 747 |
| 4 | Phypoly_transcript_04566 | *dimB* | Basic-leucine zipper transcription factor B | *Dictyostelium discoideum* | Q54ER9 | 3E-59 | 48 | 602 |
| 5 | Phypoly_transcript_16372 | *ligA* | Checkpoint protein hus1 homolog | *Dictyostelium discoideum* | Q54NC0 | 7E-61 | 41 | 271 |
| 6 | Phypoly_transcript_17606 | *ralA* | Circularly permutated Ras protein 1 | *Dictyostelium discoideum* | Q75J93 | 7E-21 | 39 | 842 |
| 7 | Phypoly_transcript_12276 | *cycU4A* | Cyclin-U4-1 | *Arabidopsis thaliana* | O80513 | 4E-22 | 36 | 202 |
| 8 | Phypoly_transcript_09630 | *cycU4B* | Cyclin-U4-3 | *Arabidopsis thaliana* | Q9FKF6 | 1E-21 | 34 | 219 |
| 9 | Phypoly_transcript_05820 | *ccnY2* | Cyclin-Y | *Mus musculus* | Q8BGU5 | 3E-39 | 37 | 341 |
| 10 | Phypoly_transcript_00893 | *msh5* | DNA mismatch repair protein MSH5 | *Arabidopsis thaliana* | F4JEP5 | 3E-147 | 36 | 807 |
| 11 | Phypoly_transcript_08670 | *ncbP* | Eukaryotic translation initiation factor 4E type 2 | *Homo sapiens* | O60573 | 4E-36 | 35 | 245 |
| 12 | Phypoly_transcript_00491 | *dhkH* | Hybrid signal transduction histidine kinase J | *Dictyostelium discoideum* | Q54YZ9 | 6E-57 | 42 | 2062 |
| 13 | Phypoly_transcript_02044 | *mkkA* | Mitogen-activated protein kinase kinase kinase A | *Dictyostelium discoideum* | Q54R82 | 2E-109 | 59 | 942 |
| 14 | Phypoly_transcript_04470 | *morC* | MORC family CW-type zinc finger protein 3 | *Mus musculus* | F7BJB9 | 1E-121 | 52 | 942 |
| 15 | Phypoly_transcript_03884 | *msh4* | MutS protein homolog 4 | *Homo sapiens* | O15457 | 1E-157 | 36 | 936 |
| 16 | Phypoly_transcript_00429 | *znfX1* | NFX1-type zinc finger-containing protein 1 | *Mus musculus* | Q8R151 | 4E-149 | 40 | 1909 |
| 17 | Phypoly_transcript_02552 | *pldA* | Phosphatidylinositol-glycan-specific phospholipase D | *Rattus norvegicus* | Q8R2H5 | 2E-71 | 32 | 843 |
| 18 | Phypoly_transcript_04506 | *pldB* | Phosphatidylinositol-glycan-specific phospholipase D | *Homo sapiens* | P80108 | 5E-49 | 29 | 840 |
| 19 | Phypoly_transcript_02197 | *pldC* | Phospholipase D1 | *Schizosaccharomyces pombe* | Q09706 | 8E-55 | 31 | 1369 |
| 20 | Phypoly_transcript_11692 | *pwiA* | Piwi-like protein Ago3 | *Bombyx mori* | A9ZSZ2 | 8E-37 | 31 | 926 |
| 21 | Phypoly_transcript_06819 | *agdA* | Probable ADP-ribosylation factor GTPase-activating protein AGD14 | *Arabidopsis thaliana* | Q8RXE7 | 2E-17 | 35 | 649 |
| 22 | Phypoly_transcript_06861 | *bzpJ* | Probable basic-leucine zipper transcription factor J | *Dictyostelium discoideum* | Q554P0 | 6E-25 | 51 | 787 |
| 23 | Phypoly_transcript_08298 | *pptB* | Probable protein phosphatase 2C 34 | *Oryza sativa subsp. japonica* | Q94H98 | 1E-28 | 30 | 380 |
| 24 | Phypoly_transcript_00697 | *meiB* | Protein MEI2-like 5 | *Arabidopsis thaliana* | Q8VWF5 | 7E-80 | 74 | 800 |
| 25 | Phypoly_transcript_11265 | *ml2* | Protein MEI2-like 5 | *Arabidopsis thaliana* | Q8VWF5 | 4E-27 | 35 | 800 |
| 26 | Phypoly_transcript_00670 | *pumA* | Pumilio homolog 1 | *Arabidopsis thaliana* | Q9ZW07 | 2E-138 | 61 | 968 |
| 27 | Phypoly_transcript_03260 | *uchA* | Secretory immunoglobulin A-binding protein EsiB | *Escherichia coli O6:H1* | A0A0H2VDN9 | 1E-13 | 34 | 490 |
| 28 | Phypoly_transcript_13976 | *hcaD* | Testis-expressed protein 30 | *Mus musculus* | Q3TUU5 | 5E-11 | 34 | 225 |
| 29 | Phypoly_transcript_16748 | *gapA* | Uncharacterized protein C622.14 | *Schizosaccharomyces pombe* | O94601 | 1E-24 | 44 | 321 |
| 30 | Phypoly_transcript_06284 | *cudA* | Uncharacterized protein DDB_G0286447 | *Dictyostelium discoideum* | Q54LR3 | 7E-51 | 47 | 521 |

Using the pseudo-cell time series as a control shows that the temporal order of switching events leading to the onset of an extended state in our dataset became apparent only through the use of true longitudinal time series of individual cells.

#### Petri net representation of the temporal order of Extended States

The iGraph plots in SI Fig. 14 show the global temporal order in which the analyzed genes enter an extended state. To obtain information about the behavior of individual cells, a separate Petri net was constructed for each unbranched path in the iGraph plot. SI Fig. 15A shows an example for the 4-plet (gene set) *bzpJ, dimB, msh5*, and *msh4*. As can be seen, the 16 cells start in 13 different states and gradually converge toward the extended states of these four genes. Thirteen cells converge into a state represented by the place named *Xon*_*Xoff*_*Xon*_*Xon*, which designates the individual states of the aforementioned genes in that order (see Fig. 1E,F). Many parallel paths through the Petri net are taken to reach this place, demonstrating a high degree of concurrency in the population as the cells enter the extended states. These parallel paths result from the combinatorics of the various states in which the four genes reside before or after entering an extended state. Accordingly, the degree of concurrency decreases over time. To observe how the cell population behaves focussed on genes entering extended states, Petri nets were constructed based on a *Xonly Node Table*. To obtain the *Xon*ly-Node-Table, all states that are not *Xon* or *Xoff* were replaced by *na*. In the resulting Petri net (SI Fig. 15B), the sequential entry of the four genes into the extended state can easily be traced, as many of the concurrent paths in SI Fig. 15A merge into one. The result is a cascade-like structure followed by most cells, with deviations from the main paths being easily recognizable.

**SI Figure 16.**
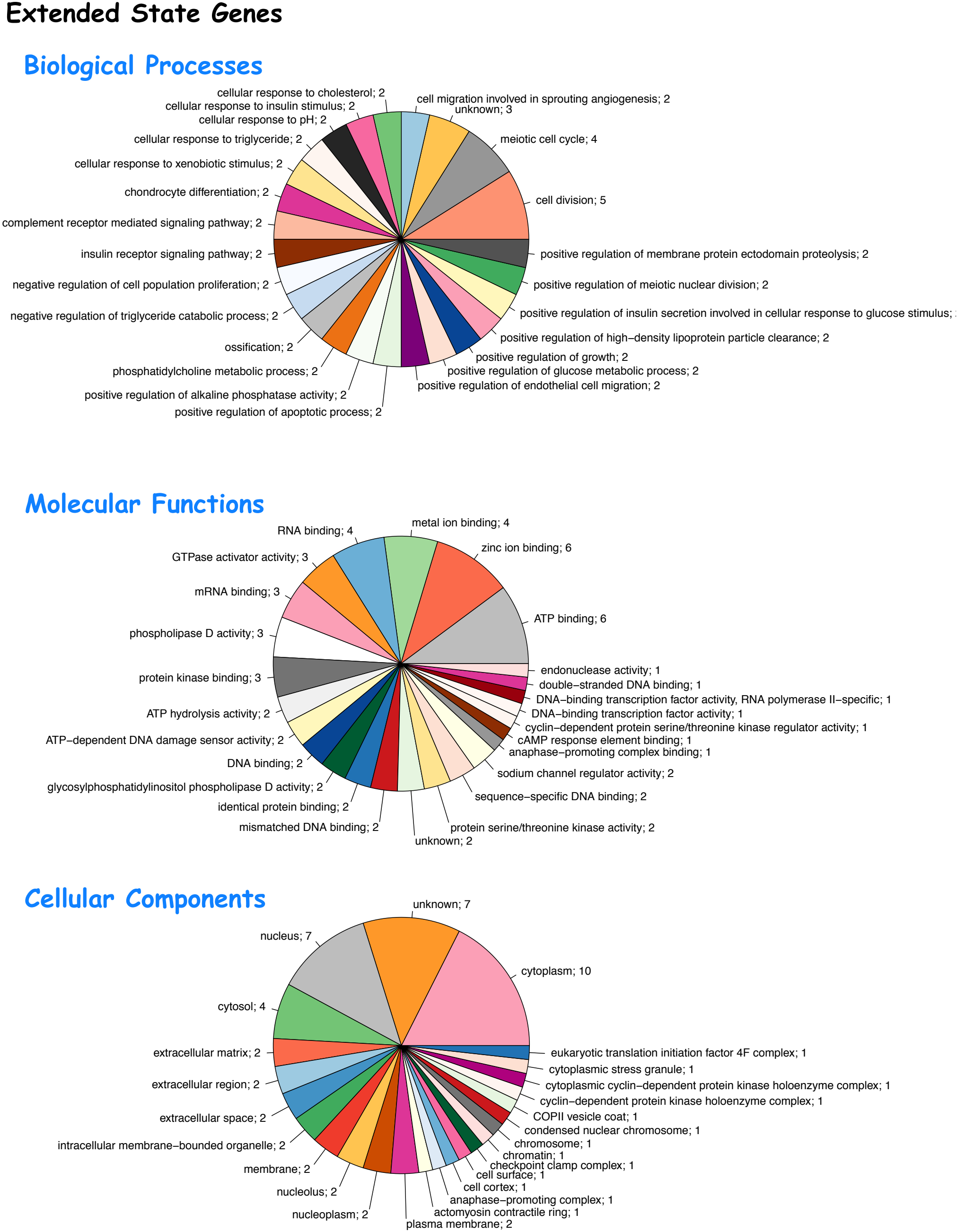
Gene Ontology terms of the closest homologs of genes with a significant temporal order of entry into an extended state. The graph shows the frequency distribution of the 25 most frequent biological function terms for the genes in the dataset that entered an extended state (*Xon* or *Xoff*). The most frequent superterms are regulation, response, differentiation, signaling, and cell cycle and proliferation control.

## Appendix 1 About Hubs and Diamonds - formal Definitions

### Definition 1

**(Petri net, Syntax)**. *A* Petri net *is a quadruple N* = (*P, T, f, m*_0_), *where*

- *P and T are finite, nonempty, and disjoint sets. P is the set of* places *(in the figures represented by circles). T is the set of* transitions *(in the figures represented by rectangles)*.
- *f* : ((*P* × *T*) [ (*T* × *P*)) → ℕ_0_ *defines the set of directed* arcs, *weighted by nonnegative integer values*.
- *m*_0_ : *P* → ℕ_0_ *gives the* initial marking.

We introduce the following notations.

The **preset** of a node *x* ∈ *P* ⋃ *T* is defined as •*x* := {*y* ∈ *P* ⋃ *T*|*f* (*y, x*) ≠ 0}, and its **postset** as *x*• := {*y* ∈ *P* ⋃ *T*|*f* (*x, y*) ≠ 0}. Altogether we get four types of sets:

- •*t*, the set of immediate preplaces of a transition *t*,
- *t*•, the set of the immediate postplaces of a transition *t*,
- •*p*, the set of the immediate pretransitions of a place *p*,
- *p*•, the set of the immediate posttransitions of a place *p*.

By generalisation, the preset for a set of nodes *X* ⊆ *P* ⋃ *T* is defined as the set of all prenodes • *X* := ⋃ _*x*∈*X*_ •*x*, and the set of all postnodes as *X*• := ⋃_*x*∈*X*_ *x*•.

### Definition 2

**(State Machine)**. *A Petri net is called* State Machine *(SM), if* ∀*t* ∈ *T*: |• *t*| = |*t*• | ≤ 1.

*Remarks:*

- In a state machine, there are neither forward branching nor backward branching transitions.
- If there are no boundary transitions, it holds: ∀*t* ∈ *T* : |• *t* | = |*t*•| = 1.

### Definition 3

**(Parallel transitions)**. *Two transitions* t_1_ *and* t_2_ *are called* parallel transitions, *if* • *t*_1_ = • *t*_2_ *^ t*_1_• = *t*_2_•.

*By generalisation, a set of transitions X*_*T*_⊆ *T are called parallel transitions, if* ∀*t*_1_, *t*_2_ ∈ *X*_*T*_ : • *t*_1_ = • *t*_2_ *^ t*_1_• = *t*_2_•

*Remark:*

- If *N* is a state machine without boundary transitions, the cardinality of the pre- and postsets of parallel transitions is always one; i.e. it holds additionally |• *X*_*T*_ | = |*X*_*T*_ • | = 1.

### Definition 4

**(Hub)**. *Let τ be a user-defined threshold and N be a state machine. Then a place p* ∈ *P with* |• *p*| ≥ *τ is called a hub place, if* |• (• *p*)| ≥ 2.

*In words:* a hub is formed by a place *p* with at least *τ* ingoing arcs, emerging from upstream transitions that are directly connected to at least two different preplaces, with *τ* being a user-defined threshold.

*Remarks:*

- Typically, the upstream transitions have pairwise different preplaces. However, the upstream transitions may also share preplaces, but there have to be at least two different preplaces.
- As a special case, a hub place may represent a terminal state (without any outgoing arcs).
- By definition, an initial state, in which many cells start, is not considered as a hub place, because the preceding states are not known.

### Definition 5

**(Diamond)**. *Let τ be a user-defined threshold and N be a state machine. Then two places p*_*in*_, *p*_*out*_ ∈ *P form a* diamond, *if* |*p*_*in*_• ⋂ • *p*_*out*_| ≥ *τ*.

*In words:* a **diamond** is formed by two places, called input and output place, which are directly connected by at least *τ* parallel transitions, with *τ* being a user-defined threshold.

*Remarks:*

- Besides these connecting parallel transitions, the input place may have further posttransitions, and the output place may have further pretransitions.
- The input place of a diamond is always a hub place, if it does not represent an initial state or is not at the same time the output place of another diamond.
- The output place may also be a hub, if it receives input from additional ingoing arcs with transitions directly connected to preplaces other than the input place of the diamond.

## Appendix 2

**SI Figure A2-1.**
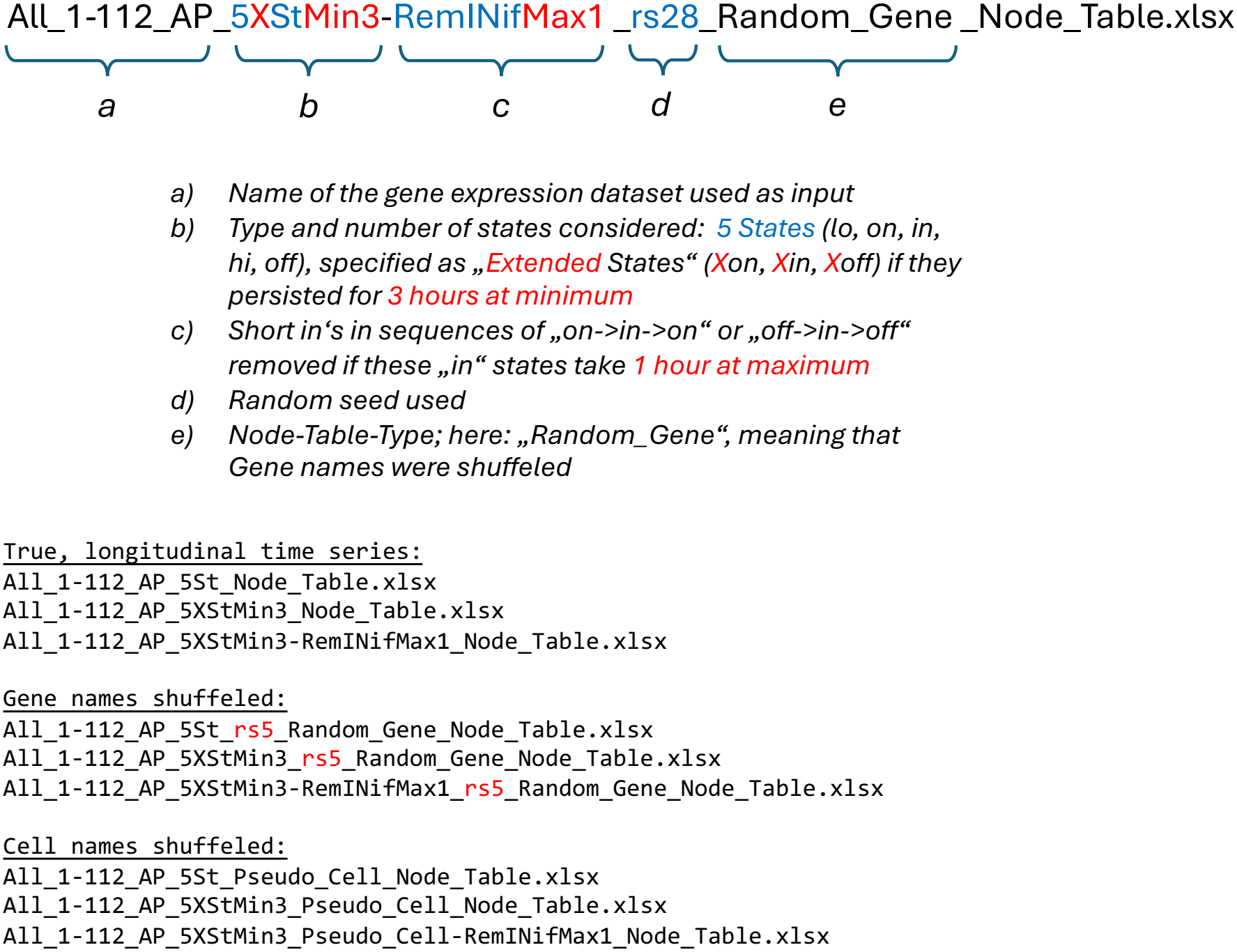
Naming scheme for the Node Tables used in this study. The diagram illustrates how naming elements are combined to name the files containing the corresponding Node Tables. Depending on the type of a Node Table (see diagram SI Fig. 7), the name may consist of several elements: (a) the name of the gene expression dataset used to generate the Node Table; (b) the type and number of states considered. For example, the abbreviation “5XStMin3” indicates that five states (*lo, on, in, hi*, and *off*) and their corresponding extended states (*Xon, Xin*, and *Xoff*) were considered, where states that persisted for at least three hours were defined as extended states, *X*. The expression in (c) indicates that short *in* states within sequences of *on*->*in*->*on* or *off*->*in*->*off* were removed (i.e., converted to *on* or *off*, respectively) if these *in* states lasted no more than one hour (where Max1 denotes one hour or less). (d) gives the value of the random seed used to obtain the specific result, where applicable. The expression in (e) indicates that this is a Random Gene Node Table (see Computational Methods in this Supplementary). The lower part of the Figure displays a list of explicit names of Node Tables used in this study.

## Use of AI for code generation

GTP-4 and GTP-5 (versions accessed in August and September 2025) were use to develop the R code to generate the Dumbbell plots, to perform the statistical analysis of genes entering an extended state in a pairwise temporal order, and to create the graphical display of the statistical analysis results (part of Figure 4 and Supplementary Information Figures 8, 10–13). All AI-generated source code is labelled as such (see ReadMe for module 3-1 of the computational pipeline (Marwan et al. 2026)). It was thoroughly tested for functionality, and manually adapted to the project’s requirements. The source code of the function performing the statistical analysis was verified line by line to ensure correct performance. WM, one of the authors of this publication, takes full responsibility for ensuring that the final version of the code functions correctly.

